# The RBPome of influenza A virus mRNA reveals a role for TDP-43 in viral replication

**DOI:** 10.1101/2023.03.21.533609

**Authors:** Maud Dupont, Tim Krischuns, Quentin Giai-Gianetto, Sylvain Paisant, Stefano Bonazza, Jean-Baptiste Brault, Thibaut Douché, Joel I Perez-Perri, Matthias W Hentze, Stephen Cusack, Mariette Matondo, Catherine Isel, David G Courtney, Nadia Naffakh

## Abstract

Recent technical advances have significantly improved our understanding of the RNA-binding protein (RBP) repertoire present within eukaryotic cells, with a particular focus on the RBPs that interact with cellular polyadenylated mRNAs. However, recent studies utilising the same technologies have begun to tease apart the RBP interactome of viral mRNAs, notably SARS-CoV-2, revealing both similarities and differences between the RBP profiles of viral and cellular mRNAs. Herein, we comprehensively identified the RBPs that associate with the NP mRNA of an influenza A virus. Moreover, we provide evidence that the viral polymerase is essential for the recruitment of RPBs to viral mRNAs through direct polymerase-RBP interactions during transcription. We show that loss of TDP-43, which associates with the viral mRNAs, results in lower levels of viral mRNAs within infected cells, and a decreased yield of infectious viral particles. Overall, our results uncover an important role for TDP-43 in the influenza A virus replication cycle via a direct interaction with viral mRNAs, and point to a role of the viral polymerase in orchestrating the assembly of viral mRNPs.

## INTRODUCTION

Influenza A viruses (IAVs) are major human and animal pathogens (1). Infectious virions contain eight viral RNA (vRNA) segments of negative polarity, which together encode ten major and several auxiliary viral proteins. Each vRNA is encapsidated with the nucleoprotein (NP) and one copy of the viral RNA-dependent-RNA polymerase (FluPol), to form macromolecular complexes called viral ribonucleoproteins (vRNPs). Unlike most RNA viruses, IAVs transcribe and replicate their genome in the nucleus of the host cell. The FluPol transcribes mRNA from vRNA templates, and performs genome replication by the synthesis of a complementary full-length RNAs (cRNAs) which in turn serve as a template for the synthesis of vRNAs (2, 3). A subset of viral mRNAs are exported to the cytoplasm via the NXF1-NXT1 pathway to be translated into proteins by the host cell translation machinery. However, for some transcripts, notably the early unspliced viral mRNAs, the mechanisms for nuclear export remains unclear (4). Newly synthesized vRNP components are re-imported into the host cell nucleus and are assembled together with replicated genomic vRNAs into *de novo* vRNPs. Late during infection, vRNPs are exported out of the nucleus via the CRM1 export pathway, transported to the plasma membrane and incorporated into progeny virions.

Influenza mRNAs are 5’-capped and 3’-polyadenylated, like cellular mRNAs. However, they are synthesized by the FluPol through a very particular process. Initiation of viral mRNA trancription occurs via a mechanism known as “cap-snatching” whereby the capped 5’ extremities of nascent host RNA polymerase II (RNAPII) transcripts are cleaved by the FluPol and subsequently used as primers to initiate transcription (5, 6). Polyadenylation and termination occur at a stretch of five to seven U residues near the 5’-vRNA end, where stuttering of the FluPol occurs and results in the formation of a 3’-poly(A) tail (3). Unlike cellular pre-mRNAs, most of the viral mRNAs are intronless. However, three of the eight vRNA segments, the M, NS and PB2 segments, produce both spliced and unspliced mRNAs, thereby expanding the coding capacity of the viral genome.

In eukaryotic cells, various proteins become associated to mRNAs as they are being transcribed and processed to form mRNA-protein complexes (mRNPs). The associated proteins control the fate of an mRNA, including splicing, polyadenylation, nuclear export, localization in the cytoplasm, translation efficiency, quality control, stability and decay and the composition of mRNPs varies depending on the nature and the life cycle step of the mRNA (7, 8) . Even though the landscape of proteins associated to cellular mRNAs has been explored on a global level (9, 10), little is known about the interactome of specific cellular mRNAs (11). Influenza viruses are co-opting the host transcriptional, splicing and nuclear export machineries to ensure efficient viral gene expression. The FluPol is actively recruited to the sites of active RNAPII transcription, and a direct interaction between the FluPol and the RNAPII C-terminal domain was shown to be essential for cap-snatching (6, 12, 13). Mapping of virus-host protein-protein interactions and genetic screens have led to the identification of numerous transcription, splicing and mRNA export factors that are potentially relevant to IAV gene expression, only a small fraction of which has been thoroughly investigated (reviewed in (5)). To date, very little is known about the proteins that are assembled with the viral mRNAs to form viral messenger ribonucleoproteins (mRNPs), and how these RNA-binding proteins (RBPs) control the fate of spliced and unspliced viral mRNAs and their delivery to the cellular translational machinery in the cytoplasm.

In this study, we focused on the highly-abundant, intronless viral mRNA encoding the NP protein. We applied a powerful combination of protein-RNA cross-linking and RNA interactome capture using a sequence-specific antisense RNA probe (14), and we identified 51 proteins that directly bind to the NP mRNA in IAV-infected human cells. By combining our proteomics data with loss-of-function experiments, we uncovered cellular factors important for a productive viral infection, among which the host protein TDP-43 was shown to be recruited to the transcribing viral polymerase by an RNA-independent interaction between FluPol and TDP-43. Overall, our data provide the first insights into influenza mRNP composition and function, and reveal an essential role of the transcribing viral polymerase in recruiting cellular RBPs to viral mRNAs.

## MATERIALS AND METHODS

### Cells and viruses

A549 (ATCC CCL-185) and HEK-293T cells (ATCC CRL-3216) were grown in complete Dulbecco’s modified Eagle’s medium (DMEM, Gibco) supplemented with 10% (v:v) fetal calf serum (FCS) and 1% penicillin-streptomycin. Madin-Darby Canine Kidney (MDCK) and MDCK-SIAT (15) cells, kindly provided by the National Reference Center for Respiratory Viruses (Institut Pasteur, Paris, France) and by Dr. M. Matrosovich (Philipps Universität, Marburg, Germany) respectively, were grown in Modified Eagle’s Medium (MEM) supplemented with 5% FCS, 100 U/mL penicillin and 100 µg/mL streptomycin. STING-37 cells expressing the Firefly luciferase under the control of Interferon-Stimulated Response Elements (16), kindly provided by Dr. P.O. Vidalain (CIRI, Lyon), were grown in DMEM supplemented with 5% FCS and 400 µg/mL geneticin. The recombinant viruses A/WSN/33 (WSN), WSN-PB2-2A-Nanoluc, WSN-PB2-Gluc1 and WSN-PB2-2A-mCherry were produced by reverse genetics as described in (17, 18). The human seasonal influenza viruses A/Bretagne/7609/2009 (H1N1pmd09) and A/Centre/1003/2012 (H3N2) serially passaged on A549 cells (19) were kindly provided by Dr. C. Demeret (Institut Pasteur, Paris, France).

### RNA interactome capture (eRIC)

The eRIC protocol used here was essentially as previously published (14), with the exception that an NP-mRNA-specific capture probe bound to magnetic beads was used in place of a 20 thymidine nucloetides (dT) probe. This is due to the fact that we aimed to specifically capture NP mRNAs from infected cells, while the original eRIC method was designed to capture all mRNAs indiscriminately by binding to their poly(A) tails. Similar to the dT probe in the original method, our probe alternated between standard nucleic acids and locked nucleic acids (LNA, indicated by a + preceding the modified nucleotide in the following sequence of the NP probe: [+g]aa[+g]ct[+t]ga[+t]ac[+t]ct[+t]ag[+a]tc[+t]), and was conjugated to carboxy beads.

For each condition nine 150 cm^2^ plates of A549 cells were grown to 80% confluency. Cells were then washed with PBS and either infected with WSN at a multiplicity of infection (MOI) of 5 PFU/cell, or not infected (mock control). Cells were further incubated for 8 h, before media was removed and cells washed with ice cold PBS. Cells were then covered with a minimal layer of ice-cold PBS and either UV crosslinked at 254 nm (AnalyticJena Crosslinker), or left untreated (non-crosslinked control). Cells were then scraped from the dishes in PBS and then the cells were pelleted at 2500 g for 10 min at 4°C. After removal of the PBS from the pellet, cells were resuspended in 15 mL of ice cold lysis buffer (20 mM Tris-HCl pH 7.5, 500 mM LiCl, 1 mM EDTA, 5 mM DTT and 0.5% LiDS) and incubated on ice for 30 min before being homogenised by serial passage through a 22-gauge syringe followed by a 27-gauge syringe. The lysate was then snap frozen and stored at -80°C.

The frozen lysate was thawed at 37°C before the addition of RNase inhibitor and incubation at 55°C for 10 min. The lysate was then placed on ice for 5 mn before being clarified at 8000 g for 10 min at 4°C. The supernatant was incubated with 300 μl of magnetic beads conjugated to the LNA-NP probe at room temperature for 1 h before beads, now with bound NP mRNA, were captured on a magnet and lysate was removed. Beads were then washed twice in one volume of pre-warmed (37°C) lysis buffer and rocked for 5 min prior to three successive washes with one volume of each of the pre-warmed wash buffers #1 (20 mM Tris-HCl pH 7.5, 500 mM LiCl, 1 mM EDTA, 5 mM DTT and 0.1% LiDS), #2 (20 mM Tris-HCl pH 7.5, 500 mM LiCl, 1 mM EDTA, 5 mM DTT and 0.02% NP40) and #3 (20 mM Tris-HCl pH 7.5, 200 mM LiCl, 1 mM EDTA, 5 mM DTT and 0.02% NP40). Beads were then resuspended in 150 μl of nuclease-free water. A 15 μl aliquot was removed and bound RNA was heat-eluted at 95°C for 10 min and analysed using an Agilent 2100 Bioanalyser. The remaining 135 μl water-bead mix was incubated with 15 μl of RNase cocktail (0.2 μl RNase A/T1 (ThermoFisher, AM2288) in 100 mM Tris-HCl pH 7.5) at 37°C for 30 min. Beads were captured and the supernatant (containing the NP mRNA interacting proteins) was collected and stored at -80°C for mass spectrometry analysis.

### Tandem Mass Tag (TMT)-based quantitative mass spectrometry

The eRIC elution samples were treated using the SP3 protocol to reduce and alkylate the proteins and to remove any chemicals as described previously (14, 20). Proteins were digested using sequencing grade modified trypsin (Promega) with a ratio of 1:50 (enzyme:protein) at 37°C overnight. Digestion was stopped with formic acid (FA) at a final concentration of 1% and the peptides were desalted using the Stage-Tips method as described in (21, 22). Briefly, two C18 Empore discs were stacked on a P200 tip before two priming steps with methanol and with ACN 80%, FA 0.1%. Membranes were washed with FA 0.1% before the sample were loaded. Finally, three washing steps were performed with 1% FA before an elution with ACN 80%, FA 0.1%. All samples were dried and resuspended in HEPES 100 mM, pH 8.5.

For TMT labeling, TMT10plex (90110 – Thermo Scientific) was resuspended with anhydrous ACN and added to each sample with a 8:1 ratio (TMT reagent:peptide). Following a 1 h incubation at room temperature, the TMT was quenched with hydroxylamine at a final concentration of 5% during 15 min at room temperature. Labeled samples were pooled and freeze-dried before resuspension in FA 1%. Samples were desalted using the Stage-Tips method described above.

Data were acquired using a nanochromatographic system (Proxeon EASY-nLC 1200 – Thermo Scientific) coupled on-line to a Q Exactive^TM^ HF mass spectrometer. For each sample, the peptides in solution were injected into a reverse phase column (home-made column, 34cm x 75 µm ID, 1.9 µm particles, 100 Å pore size, ReproSil-Pur Basic C18 – Dr. Maisch GmbH) after an equilibration step in solvent A (H_2_O, 0.1% FA). Peptides were eluted with a multi-step gradient from 2 to 7% of buffer B (ACN 80%, FA 0.1%) during 5 min, 7 to 23% during 70 mn, 23 to 45% during 30 min and 45 to 95% during 5 min at a flow rate of 250 nL/mn for up to 132 mn. The column temperature was set to 60°C. Mass spectra were acquired using the Xcalibur software using a data-dependent Top 10 method with a survey scan (300-1700 m/z) at a resolution of 120,000 and MS/MS scans (fixed first mass 100 m/z) at a resolution of 60,000. The AGC target and maximum injection time for the survey scans and the MS/MS scans were set to 3×10^6^, 50ms and 10^5^, 100ms respectively. The isolation window was set to 1.2 m/z and normalized collision energy fixed to 28 for HCD fragmentation. We used a minimum AGC target of 2×10^3^ for an intensity threshold of 2×10^4^. Unassigned precursor ion charge states as well as 1, 8 and >8 charged states were rejected and peptide match was preferred. Exclude isotopes was enabled and selected ions were dynamically excluded for 30 sec.

### Mass spectrometry data analysis

Raw data were analysed using MaxQuant software version 1.6.10.43 (23) using the Andromeda search engine (24). The MS/MS spectra were searched against multi databases: a UniProt Homo sapiens database (20,415 entries the 19/10/2019), a Influenza A virus database (11 entries the 07/01/2020) and the Bovin pancreatic ribonuclease (UniProt ID: P61823). Usual known mass spectrometry contaminants and reversed sequences of all entries were included. Andromeda search was performed choosing trypsin as specific enzyme with a maximum number of two missed cleavages. Possible modifications included carbamidomethylation (Cys, fixed), oxidation (Met, variable), Nter acetylation (variable). The mass tolerance in MS was set to 20 ppm for the first search then 4.5 ppm for the main search and 20 ppm for the MS/MS. Maximum peptide charge was set to seven and seven amino acids were required as minimum peptide length. The “match between runs” feature was applied with a maximal retention time window of 0.7 min. One unique peptide to the protein group was required for the protein identification. A false discovery rate (FDR) cutoff of 1 % was applied at the peptide and protein levels. With the TMT labeling, reporter ion MS2 signals were corrected for isotope impurities and used for the statistical analysis. The mass spectrometry proteomics data have been deposited to the ProteomeXchange Consortium via the PRIDE partner repository (25) with the dataset identifier PXD040783.

To find the proteins more abundant in one condition than in another, the intensities quantified using Maxquant were compared. Reverse hits, potential contaminants, and proteins not well identified with a 1% FDR (“Only identified by site”) were first removed from the analysis. Only proteins identified with at least one peptide that is not common to other proteins in the FASTA file used for the identification (at least one “unique” peptide) were kept. Additionally, only proteins with at least six quantified intensity values in one of the two compared conditions were kept for further statistics. After this filtering, intensities of the remaining proteins were first log-transformed (log2). Next, intensity values were normalized by median centering within conditions (section 3.5 in (26)). Missing values were imputed using the impute.mle function of the R package imp4p (27). Statistical testing was conducted using a limma t-test thanks to the R package limma (28). An adaptive Benjamini-Hochberg procedure was applied on the resulting p-values thanks to the function adjust.p of the cp4p R package (29) using the robust method described in (30) to estimate the proportion of true null hypotheses among the set of statistical tests. The proteins significantly more abundant in the *infected crosslinked* condition than in both controls (*non infected-crosslinked* and *infected-non crosslinked*) were defined as representing the “Core interactome” (adjusted p-value lower than a false discovery rate level (FDR) of 1% and log2(fold-change) higher than 2) and the “Expanded interactome” (adjusted p-value lower than a FDR of 10% and log2(fold-change) higher than 1.5).

Protein-protein interaction networks have been determined using the app stringApp (31) of Cytoscape (32). Widths of the edges are function of the “experiments” scores provided by STRING (33). This score reflects the confidence we can have in an interaction from known interactions listed in several experimental databases. Colors are function of the measured log2(fold-change). Enrichment analyses of the proteins of interest were also performed using stringApp. All the proteins identified by MS have been used as background for the enrichment tests. A significantly low p-value means the proportion of proteins related to a term is significantly superior in the considered interactome than in this background.

### siRNA-based assays

Small interfering RNAs (siRNAs) (Dharmacon ON-TARGETplus SMARTpools, individual siRNAs and Non-targeting Control pool) were purchased from Horizon Discovery. A549 cells seeded in 96-well plates (10^4^ cells/well) were transfected with 37.5 nM of siRNA, using 0.3 µL of the DharmaFECT1 transfection reagent (Horizon Discovery), and infected at 48 hours post-infection (hpt) with the WSN-PB2-2A-Nanoluc virus at a MOI of 10^-3^ PFU/cell. Luciferase activity was measured at 24 hours post-infection (hpi) using the Nanoluc Luciferase kit (Promega) and a Centro XS3 luminometer (Berthold). Cell viability in the presence of siRNAs was assessed in the same conditions, using the CellTiter-Glo Luminescent Viability Assay kit (Promega).

### Plasmids

Plasmids pSPICA-Gluc2-TOP3B, pcDNA3.1-Gluc2-RBP1, pCI-WSN-PB1, pCI-WSN-PA, pCI-WSN-PB2-Gluc1, pCI-WSN-PB1-Gluc1, pCI-Gluc1, pCI-Gluc2 and pCI-PB1-3xFlag were described previously (13, 34, 35). The TDP-43, hnRNPA2B1, hnRNPH1, SF3B4 and SF3A3 open reading frames (ORFs) were obtained from the Human ORFeome resource (hORFeome v3.1) and were subcloned instead of the WSN-PB1 sequence in the pCI-WSN-PB1-Gluc2 or the pCI-WSN-PB1-3xFlag plasmid. The pCI-mcherry-NLS-3xFlag, pCI-chANP32A-Gluc2, pCI-ANP32A-Gluc2, and pCI-NP-Gluc1 plasmids were obtained following the same strategy. The V5-tag sequence was added between the TDP-43 sequence and the Gluc2 tag in the pCI-TDP-43-Gluc2 plasmid using annealed oligo-cloning. Site-directed mutagenesis was performed as previously described (13). PCR amplicons corresponding to subdomains of TDP-43 were subcloned instead of the full-length sequence in the pCI-TDP-43-V5-Gluc2 construct. To generate lentiviral plasmids, the V5-tag sequence followed by a Stop codon was introduced upstream of the IRES in the pLVX-IRES-Neo vector (Clontech, Addgene), using annealed oligo-cloning. The wild-type and variant TDP-43 sequences were amplified by PCR and the resulting amplicons were subcloned upstream of the V5-tag sequence in the pLVX-V5-IRES-Neo lentivirus. For CLIP-qPCR experiments pcDNA-NP and pPolI-NA plasmids from the reverse genetics system for WSN (36) were used. The pcDNA-NA plasmid was generated by PCR amplifying the NA coding region from pPolI-NA and cloning it in place of the NP gene into pcDNA-NP. To produce an HA tagged TDP-43, the coding sequence of 3xHA tags was cloned using annealed oligos in place of the 3xFlag tags in pCI-TDP-43, above. Primers and plasmid sequences are available upon request.

### CLIP-qPCR

The CLIP-qPCR protocol utilised here is essentially as described in (37). Briefly, 4×10^6^ HEK-293T cells were seeded on a 10 cm dish and transfected the next day with 5 μg of a plasmid overexpressing a 3xFlag tagged candidate RBP, or mCherry employed as a negative control. At 48 hpt cells were infected with WSN at a MOI of 5 PFU/cell and incubated for 6 h. At 6 hpi, the medium was removed and cells were washed with ice cold PBS and covered with minimal ice cold PBS. Each dish underwent UV crosslinking at 254 nm (Uvitec) to covalently bind RBPs to target RNAs. Cells were then scraped, pelleted and lysed in 250 μl of NP40 lysis buffer (HEPES-KOH pH 7.5 50 mM, KCl 150 mM, EDTA pH 8.0 2 mM, NaF 1 mM and NP40 0.5%) on ice for 30 min. Lysates were then clarified and 10% was removed as an input and boiled in Laemmli buffer at 95°C for 10 min prior to western blotting. The remaining 90% of lysate was incubated with anti-Flag antibody (Merck, F1804) coupled to Protein G Dynabeads (ThermoFisher, 10004D) and incubated on a rotator for 1 h at room temperature. After 1 h of incubation, beads were captured on a magnet, washed five times with 1 mL NP40 lysis buffer and three times with PBS. A fraction (10%) of the beads was removed, resuspended in Laemmli buffer and boiled at 95°C for 10 min prior to Western blotting. The remaining 90% of beads underwent proteinase K digestion (ThermoFisher, EO0491) at 55°C for 60 min to release bound RNA, following the manufacturer’s instructions. RNA was then isolated from the digestion by TRIzol LS (ThermoFisher, 10296028) and ethanol precipitation.

For CLIP-qPCR, plasmids were transfected at the same time as the 3xFlag-RBP or -mCherry-expressing plasmids and cells were incubated for 48 h prior to CLIP. To assess the association of RBPs with RNAPII-transcribed viral transcripts, pcDNA3.1-NP or -NA expression plasmids were used. In contrast, for the association with FluPol-transcribed viral transcripts we used pcDNA3.1-PB2, -PB1, -PA and -NP expression plasmids alongside a pPolI-NA-vRNA expressing plasmid, which constituted a mini-replicon system.

Isolated RNA then underwent cDNA conversion using oligo dT or previously published primers specific to WSN transcripts (NP-mRNA: CCA GAT CGT TCG AGT CGT TTT TTT TTT TTT TTT TCT TTA ATT GTC, NP-vRNA GGC CGT CAT GGT GGC GAA TGA ATG GAC GGA GAA CAA GGA TTG C, NP-cRNA GCT AGC TTC AGC TAG GCA TC AGT AGA AAC AAG GGT ATT TTT CTT T; NA-mRNA: CCA GAT CGT TCG AGT CGT TTT TTT TTT TTT TTT TGA ACA AAC TAC (38)). Quantification of the RNA present in these cDNAs was performed by qPCR using SYBR green (ThermoFisher) with the following primer pairs: NP-mRNA: CCA GAT CGT TCG AGT CGT & CGA TCG TGC CCT CCT TTG; NP-vRNA: GGC CGT CAT GGT GGC GAAT & CTC AAT ATG AGT GCA GAC CGT GCT; NP-cRNA: GCT AGC TTC AGC TAG GCA TC & CGA TCG TGC CCT CCT TTG; NA-mRNA: CCA GAT CGT TCG AGT CGT & TGA ATA GT GAT ACT GTA GAT TGG TCT; Actin-mRNA: ATT GGC AAT GAG CGG TTC & CGT GGA TGC CAC AGG ACT). Enrichment of RNA levels was determined using the 2^-ýýCT^ method (39).

### Split luciferase-based complementation assays

The protein complementation assays (PCA) were performed as described previously, either in an infectious context (iPCA (17)) or a in transient expression context (PCA, (13)) with minor modifications. Briefly, HEK-293T cells were seeded in 96-well white opaque plates (Greiner Bio-One, 3×10^4^ cells/ well), one day before plasmid transfection with polyethylenimine (PEI, Polysciences, Inc., 3 µL of PEI at 1 mg/mL for 1 µg of DNA). Cells were transfected with each indicated Gluc1-P1 and Gluc2-P2 plasmid (total plasmid amount: 150 ng), where P1 and P2 represent proteins or protein subdomains of interest (PCA), or transfected with 100 ng of the indicated Gluc1-P1 plasmid and subsequently infected with the WSN-PB2-Gluc1 virus at a MOI of 5 PFU/cell (iPCA), or co-transfected with 25 ng of each of pCI-chANP32A-Gluc2 and pCI-WSN-NP-Gluc1 plasmids and subsequently infected with the WSN virus at a MOI of 5 PFU/cell (control for RNase A treatment). Samples substituting Gluc1-P1 by Gluc1 or Gluc2-P2 by P2 were used as background controls. The luciferase enzymatic activities were measured using the Renilla Luciferase Assay system (Promega) and a Centro XS3 luminometer (Berthold) at 6 hpi (iPCA) or 24 hpt (PCA). Normalized Luminescence Ratios (NLR) were calculated as described previously (13). When indicated, the cell lysates were split in two halves, one of which was supplemented with RNAse A (ThermoFisher, EN0531) at a final concentration of 100 µg/mL for 30 min at 37°C before luciferase activities were measured and analysed as described above.

### Co-immunoprecipitation of PA-3xFlag and TDP-43 in infected cells

For each condition, 4-8×10^6^ HEK-293T cells were cultured in 10 cm dishes for 24 h prior to transfection. Cells were transfected with either pCI-TDP-43-3xHA or a mixture of pcDNA3-mCherry-3xHA and empty vector (4:1), as a negative control for immunoprecipitation, using PEI reagent. At 24 hpt, cells were infected at a MOI of 1 PFU/cell with WSN-PA-3xFlag and incubated for 4 h. Cells were then washed with PBS and fixed for 10 min with 1% PFA. The fixative was then removed, and crosslinking was quenched with a 10 min incubation with a 1.25M glycine solution. Cells were washed three times in PBS, scraped and lysed for 40 min in 250 µl ice-cold RIPA buffer (10mM Tris-HCl pH 8.0, 1mM EDTA, 0.5mM EGTA, 1% Triton X-100, 0.1% Sodium Deoxycholate, 0.1% SDS, 140mM NaCl) supplemented with protease inihibitors on ice. Lysates were clarified and 10% was removed and labelled as input. The remaining lysate was used for immunoprecipitation. Protein G Dynabeads were associated with the monoclonal anti-Flag antibody as described above, incubating on a rotator at 4°C for 40 min, and subsequently washed three times in ice-cold PBS. The Protein G/anti-Flag coupled beads were then added to the lysates and supplemented with 2.5 µl of RNAse Cocktail (ThermoFisher, AM2286) in order to digest and remove all RNA-mediated interactions. After 2 h, the beads were captured on a magnet and washed three times with ice-cold RIPA buffer and 3 times with ice-cold PBS. Finally, the beads and input samples were resuspended in Laemmli buffer supplemented with 10% DTT and boiled for 40 min prior to western blotting.

### CRISPR gene knock-down and lentiviral gene rescue

A lentivirus encoding Spy Cas9 was used to generate a stably expressing A549 cell line. Briefly, lentiCas9-Blast (Addgene, 52962) was transfected into HEK-293T cells along with packaging plasmids pΔ8.74 (Addgene, 22036) and pMD2G (Addgene, 12259), using PEI reagent. At 72 hpt the conditionned media, now containing lentiviral particles, was passed through a 0.45 μm-pore size filter and transferred to A549 cells. At 48 h post-transduction the media was removed and replaced with blasticidin containing media (10 μg/mL) to select for transduced, and thus Cas9-expressing, cells. Cells were then single-cell cloned in 96-well plates and individual clones were assessed for Cas9 expression by western blotting using an anti-Flag antibody as described above. This allowed us to establish a parental A549-Cas9 cell line for all subsequent knockout experiments.

To knockout TDP-43 expression, we cloned oligos encoding for TDP-43 sgRNAs, taken from the GeCKO v2 library (40), into the lentiGuide-puro plasmid (Addgene, 52963). These plasmids were confirmed by Sanger sequencing before production of lentiviral particles. Lentivirus was generated in the same way as above, overlaid onto A549-Cas9 cells and selected using puromycin (2 μg/mL). Cells were single cell cloned in 96 well plates and knockouts were determined by western blotting for TDP-43.

To rescue the expression of the wild-type or variant TDP-43 proteins in TDP-43 KO cells, a series of pLVX-TDP-43-V5-IRES-Neo recombinant lentiviruses was produced. Briefly, 4×10^6^cells HEK-293T were plated in 10 cm dishes, before co-transfection the next day with 15 µg of the pLVX-TDP-43-V5-IRES-Neo plasmid along with 10 and 5 µg of the pΔ8.74 and pMD2.G pasckaging plasmids, respectively, using PEI as described above. At 72 hpt the conditionned media was passed through a 0.45μm-pore size filter and placed over TDP-43-KO2 cells. At 48 h post-transduction, cells were splitted and cultured G418 containing media (2 mg/mL) to select for a polyclonal population of transduced, TDP-43-expressing cells. The resulting sub-populations were assessed for TDP-43 expression by western blotting.

### Antibodies, immuno2blots and immunostainings

For immunoblots, total cell lysates were prepared in Laemmli buffer. Immunoblot membranes were incubated with primary antibodies directed against the viral proteins HA (Genetex GTX127294, 1:1,500), PB2 (Genetex GTX125925, 1:5,000), NS1 (kindly provided by Dr. D. Marc, INRA-Tours, France), against A/PR/8/34 virions (41), against the V5-tag (ThermoFisher, 46-0705, 1:1000), Flag-tag (Proteintech, 66008-3-ig), HA-tag (Proteintech, 66006-2-ig), Gaussia luciferase (New England Biolabs, E8023, 1:10,000), and revealed with appropriate secondary antibodies (GE Healthcare or Sigma, A9044) and the ECL 2 substrate (Pierce). The chemiluminescence signals were acquired using the Chemidoc imaging system (Biorad) and analysed with ImageLab (BioRad).

For immunostaining, 10^5^ HEK-293T cells or 7.5×10^4^ A549-derived cells were seeded on glass coverslips (13mm diameter, #1.5, Epredia, CB00130RAC20MNZ0) in 24 well-plates. The next day, HEK-293T cells were transfected with 500 ng of the indicated plasmids per well using PEI. A549-derived cells were transfected with 500 ng of poly(I:C) (a mix of high and low molecular weight, ratio 1:1, InvivoGen, tlrl-picw and tlrl-pic) using lipofectamine 2000 (ThermoFisher, 11668027). Cells were fixed with 4% paraformaldehyde (Fisher scientific, 47377) for 20 min, washed with PBS, and incubated with a permeabilization and blocking solution (PBS 1X with 0.2% Triton X-100, 0.25% fish gelatin, 5% Normal Donkey Serum and 3% Normal Goat Serum) for 20 min. Cells were incubated overnight at 4°C with a primary antibody directed against the V5-tag (ThermoFisher, 46-0705, 1:200) or against small double-stranded RNAs (J2 antibody, Merck, MABE1134, 1:100), and for 1 h at room temperature with an appropriate secondary antibody conjugated to Alexa Fluor 488 (ThermoFisher, A-11029, 1:400) and with DAPI (ThermoFisher, 62248, 1 µg/mL). Antibodies were diluted in a staining solution (PBS, 5% Normal donkey serum, 3% Normal goat serum, 0.125% fish gelatin, 0.2% Triton X-100). Coverslips were mounted on glass slide with Fluoromount-G (Invitrogen, 00-4958-02) and examined using a Leica TCS SP8 confocal microscope with HC PL APO CS2 40X (1.3) or 63X (1.4) oil objectives. The fluorescence signals were acquired with the LAS X software and analysed with Fiji (ImageJ). Metadata can be provided upon request.

### Viral replication assays

Cells were seeded in 96 well plates (3×10^4^ cells/well), in triplicates 24 h prior to infection. To assess the accumulation of viral RNA species from infected cells, cells were infected with the WSN virus at a MOI of 3 PFU/cell. Total RNA was isolated from pooled triplicates 5 hpi with RNeasy Mini columns according to the manufacturers instructions (RNeasy Kits, Qiagen) and strand-specific qPCRs were performed as described previously (38). Briefly, total RNA was reverse transcribed using primers specific for NA mRNA, cRNA and vRNA or cellular glyceraldehyde 3-phosphate dehydrogenase (GADPH) with SuperScript^TM^ III Reverse Transcriptase (Invitrogen), and quantified using SYBR-Green (Roche) with the LightCycler^R^ 480 system (Roche). RNA levels were normalized to GAPDH and analysed using the 2^-ýýCT^ as described before (39). Alternatively, cells were infected with the WSN-PB2-2A-mCherry virus at a MOI of 10^-2^ PFU/cell. The fluorescence signals were acquired using the Incucyte S3 (Essen Bioscience) every 6 h (10x objective, 5 fields per well) and were analysed using the Incucyte software. To assess the production of infectious viral particles, cells were infected with the WSN virus (3×10^-4^ PFU/cell) or the seasonal H1N1pdm09 and H3N2 viruses (3×10^-3^ PFU/cell) in OptiMEM supplemented with 0.8 µg/mL TPCK and 1% BSA. Plaque assays were performed on MDCK cells (WSN and H3N2) or MDCK-SIAT cells (H1N1pdm09) as described in (42).

### Interferon response of A549-derived cell lines

A549-derived cells (Cas9, KO2 and KO2-TDP) seeded in 24-well plates (7×10^4^ cells/well) were transfected with 500 ng of poly(I:C) as indicated above, or mock-transfected. Cells were washed with DMEM at 4 hpt and supplemented with DMEM + 10% FCS. Supernatants were collected at 8 and 24 hpt and 100 µl were transferred onto reporter STING-37 cells seeded 24 h ahead (3×10^4^ cells/well), in white opaque 96-well plates (Greiner Bio-One) pre-treated with poly-lysine (SIGMA, P8920). STING-37 cells were lysed 24 h later in 40 µl of lysis buffer (8 % glycerol (v/v), 1.5 M tris-phosphate pH 8, 0.5 mM [(Cholamidopropyl)dimethylammonium)]-1-propanesulfonate, 0.125 % (v/v) Triton ×100, 0.17 M ThioUrea, 10 mM trans-(1,2-cyclohexanedinitrilo)tetraacetic acid) for 30 min at 25°C under gentle agitation. Luciferase activity was measured following injection of the substrate (0.5 g/L D-Luciferin and 0.25 g/L coenzyme A diluted in 100 mL buffer (25 mM Tricine pH 7.8, 5 mM MgSO_4_, 0.5 mM EDTA pH 8.5 mM DTT, 0.5 mM ATP) using a Centro XS3 luminometer (Berthold).

## RESULTS

### Identification of cellular proteins bound to influenza virus NP-mRNA in human infected cells

To identify the cellular factors that interact directly with the influenza NP-mRNA during infection, we used the enhanced RNA interactome capture method (eRIC) characterised by the use of a locked nucleic acid (LNA)-modified capture probe and the use of stringent capture and washing conditions that reinforce specificity (14) (**Figure 1A**). Briefly, A549 cells were infected at a high MOI with the A/WSN/33 (WSN) virus. At 6 hpi, when high levels of NP-mRNA are expected (43), UV-crosslinking was applied to create covalent bonds between RNA molecules and their direct protein binders. Of note, neither protein-protein nor RNA-RNA crosslinking is expected to be induced by UV light. An LNA-modified probe complementary to the viral NP-mRNA was added to the cell lysates, and the capture, washes and elution steps of the eRIC method were performed as described previously (14). The proteins co-purified with the viral NP-mRNA were analysed by tandem mass spectrometry (LC-MS/MS) upon chemical labelling with tandem mass tag (TMT). Three independent biological replicates were performed. To exclude off-target hits, we applied the eRIC method to *non infected-crosslinked* and *infected-non crosslinked* control samples in parallel.

**Figure 1.**
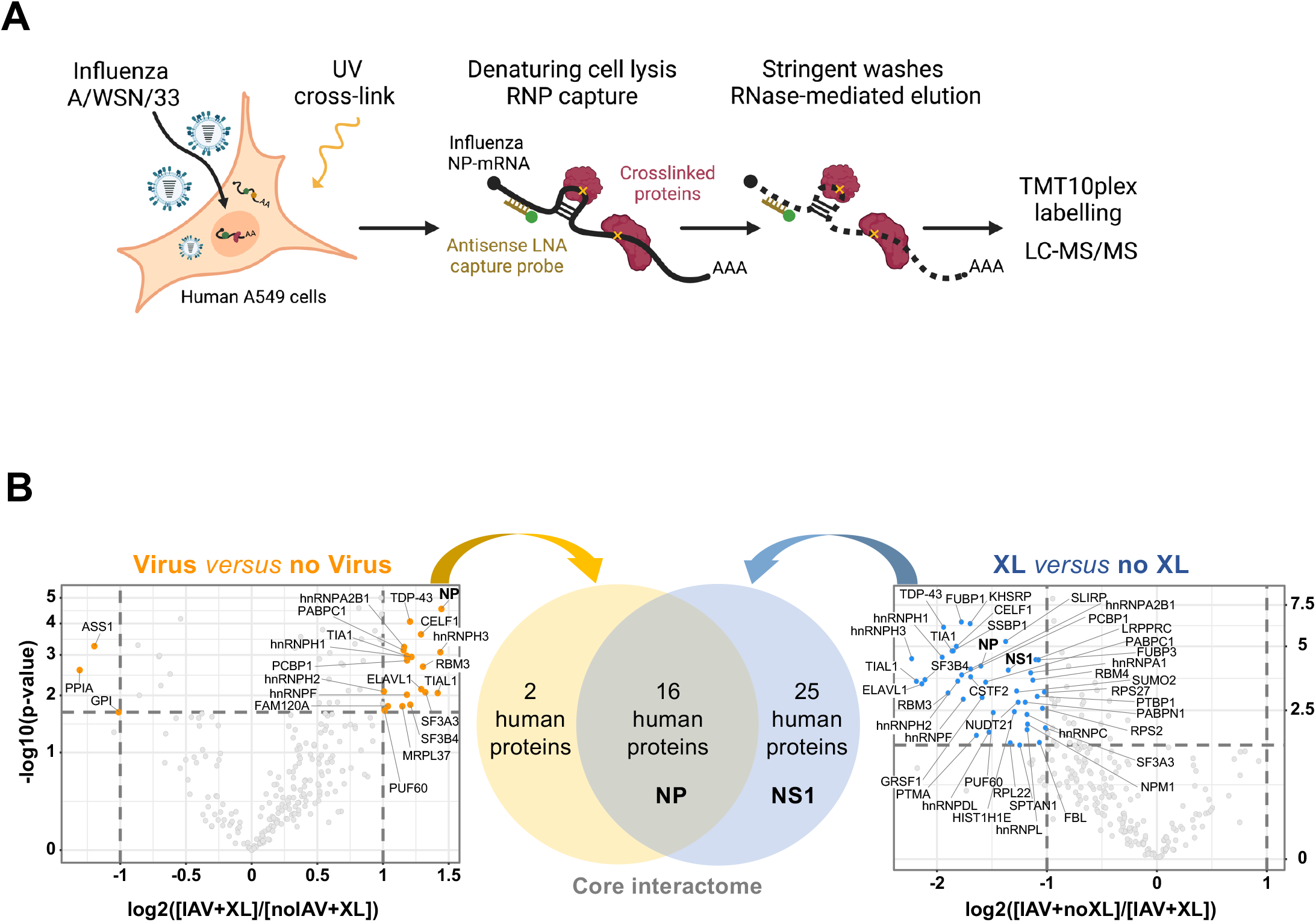
Identification of cellular proteins bound to influenza virus NP-mRNA in human infected cells. **A.** Schematic representation of the eRIC approach. **B.** Volcano plots showing the log2 fold change (x axis) and its significance (-log10(p-value) associated to a False Discovery Rate < 1%, y axis) of each protein (dots) in the eRIC experiments. Left plot: the log2 fold change refers to the enrichment in *infected crosslinked* (IAV+XL, n=3) versus *non-infected-crosslinked* (noIAV+XL, n=3) samples. Right plot: the log2 fold change refers to the enrichment in *infected crosslinked* (IAV+XL, n=3) versus *infected-non crosslinked* (IAV+noXL, n=2) samples. The set of 16 cellular proteins found to be enriched in both comparisons is referred to as the core interactome of the viral NP-mRNA. The viral NP and NS1 proteins are indicated in bold.

Correlation analysis of the mass spectrometry data showed that the independent biological replicates were highly consistent, with a Pearson’s correlation coefficients higher than 72.1%, 73.2%, and 75.2% for *infected-crosslinked* (n=3), *non infected-crosslinked* (n=3) and *infected-non crosslinked* (n=2) samples, respectively (**Supp Figure 1A-C**). Moreover, the distributions of peptide intensity values across the different samples was very similar (**Supp Figure 1D**). Using a fold-change (FC) threshold of 2 and a false discovery rate (FDR) threshold of 1%, we found 18 cellular proteins enriched in the infected over the non-infected samples, and 41 cellular proteins enriched in the crosslinked over the non crosslinked samples (**Figure 1B** and **Supp File 1**). We defined the 16 proteins enriched in both data sets as the high-confidence core interactome (**Figure 1B** and **Figure 2A**). We also defined an expanded NP-mRNA interactome with relaxed thresholds (FC > 1.5 and FDR < 10%), which includes 35 additional cellular proteins (**Figure 2B** and **Supp File 1**). Among these, the GRSF1 (44, 45), hnRNPK (46, 47), hnRNPA2B1 (48, 49), NUDT21 (50) and PABPC1 (51) proteins have already been documented as being involved in the IAV life cycle, supporting the robustness and relevance of our dataset and analysis.

**Figure 2.**
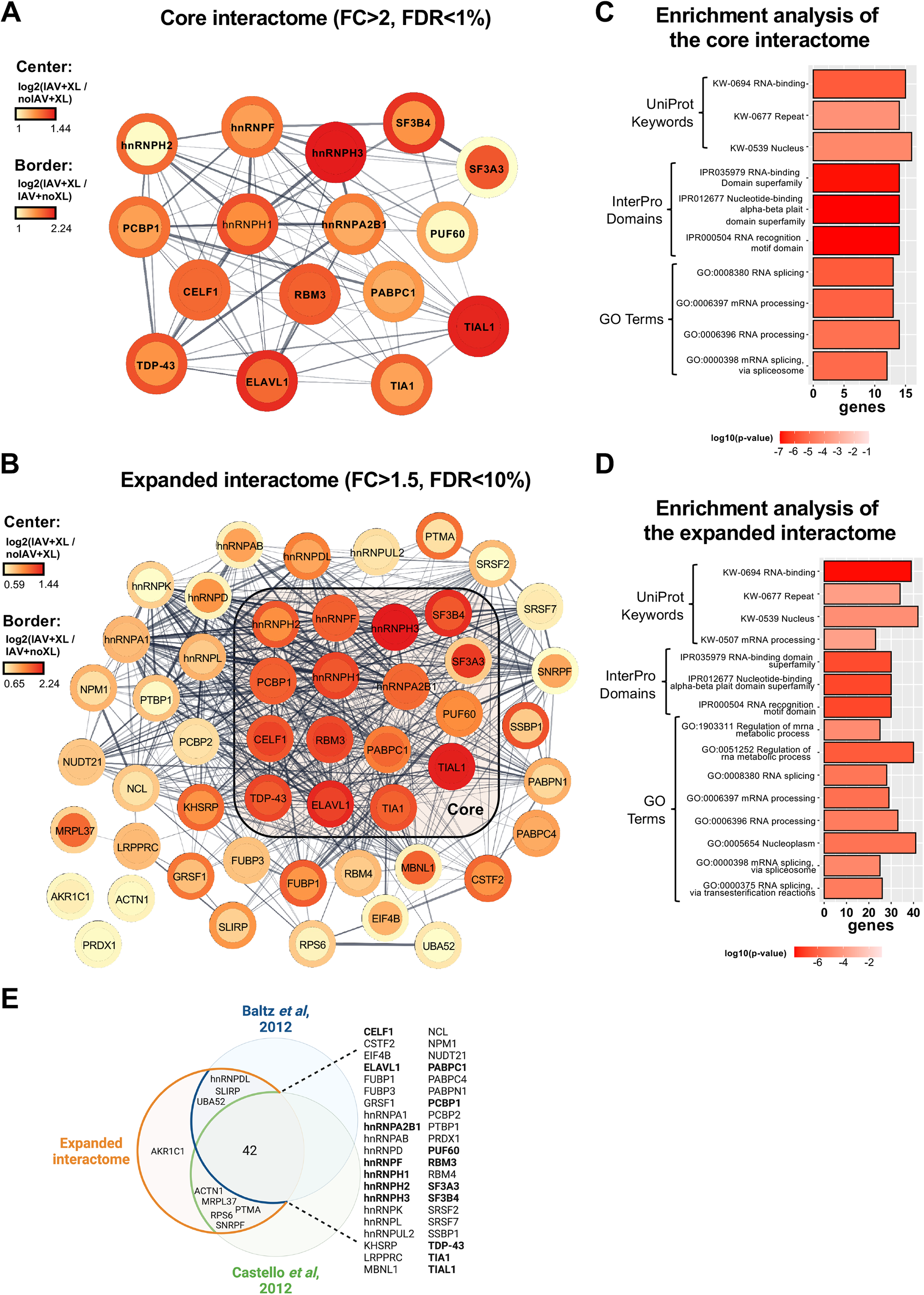
Characterisation of the NP-mRNA interactome. **A-B**. Interaction network among the 16 proteins of the core interactome (A) and the 51 proteins of the expanded interactome (B), according to the STRING database (33). Predicted protein-protein interactions are indicated by grey lines, and line thickness represents the prediction confidence score. The color scales represent the log2 fold changes characteristic of each protein and refer to the enrichment in the *infected crosslinked* versus *non-infected-crosslinked* condition (border color) or to the enrichment in *infected crosslinked* versus *infected-non crosslinked* (center color) samples. **C-D.** Gene Ontology (GO) term, InterPro Domains and UniProt Keywords enrichment analysis on the core (C) and expanded (D) interactome. The graph represents the number of genes corresponding to each indicated category (x axis) and the enrichment p-value (color scale). **E.** Venn diagram representing the overlap between the expanded interactome of the viral NP-mRNA and the cellular poly(A) mRNA interactomes published by Castello et al. (9) and Baltz et al. (10). Proteins are ranked in alphabetical order and are indicated in bold when they belong to the core interactome.

Two major viral RNA-binding proteins (RBPs), NP and NS1, were found enriched in our experiments (**Figure 1B**, FC ≥ 2 and FDR ≤ 1%). The multifunctional NS1 protein was previously shown to be co-immunoprecipitated with viral mRNAs and was proposed to play a role nuclear export and translation (4). To our knowledge there is little evidence so far for a direct binding of the NP to viral mRNAs. However, our finding is in line with the notion that NP binds to the RNA phosphate-sugar backbone with no sequence specificity (52) and is also consistent with the fact that when NP–vRNA interactions were assessed, approximately 15% of the viral sequence reads aligned to positive-sense viral RNAs (53). The viral polymerase subunits were not enriched in our experiments, in agreement with earlier observations that the viral polymerase does not remain bound to viral mRNAs after synthesis is completed (54).

### Biological functions of the viral mRNA binding proteins

We performed a Gene Ontology (GO) enrichment analysis of the core and expanded NP-mRNA interactomes, which revealed a strong enrichment for GO terms related to RNA/mRNA metabolic processes and RNA/mRNA splicing (**Figure 2C-D** and **Supp File 2**) and a strong enrichment of InterPro Domains and UniProt Keywords related to RNA-binding (e.g. IPR000504, KW-0694). The analysis also indicated that the NP-mRNA interactome was enriched in proteins with repeated domains (KW-0677) and with nucleotide-binding domains with an alpha-beta plait structure (IPR012677) (**Figure 2C-D** and **Supp File 2**). These two categories include heterogeneous nuclear ribonucleoproteins (hnRNPs), which are characterised by repeated RNA-binding domains and overall represent 31% and 24% of the core and the expanded NP-mRNA interactome, respectively. Proteins annotated in UniProt as poly(A)-binding proteins (hnRNPDL PABPC1, PABPN1, PABPC4, PABP2) or 3’UTR binding proteins (CELF1, ELAVL1, hnRNPD, hnRNPA1, hnRNPA2B1, hnRNPL, KHSRP, LRPPRC, NUDT21, PABPC1, PABPC4, TDP-43, TIA1, TIAL1) are also well-represented. Importantly, the components of the NP-mRNA interactome are strongly connected to each other in terms of protein-protein interaction (PPI), according to the STRING database (**Figure 2A-B**,grey lines, and **Supp File 2)**. Within the 51 proteins of the expanded interactome, 48 are connected to ≥ 1 protein and 16 are connected to ≥ 30 proteins (**Supp Figure 1E**).

We compared the expanded NP-mRNA interactome with two previously published datasets on the global interactome of human poly(A) mRNAs, in total about 800 proteins with an overlap of 658 common proteins (9, 10). Most of the proteins of the NP-mRNA interactome (42 out of 51) are found in both datasets while eight proteins are found in one or the other, and only one, AKR1C1, was not previously identified as a poly(A) mRNA-binding protein (**Figure 2E** and **Supp File 2**), indicating that the NP-mRNA binders correspond to a specific subset of the cellular poly(A) mRNA binders. Moreover, a large proportion of the expanded and core NP-mRNA interactomes (27 out of 51 and 10 out of 16 proteins, respectively) appear in the list of ∼200 proteins previously identified in the viral RNA-interactome of positive-stranded RNA viruses from three distinct families (55), suggesting that similar RNA biogenesis pathways are hijacked by divergent viruses (**Supp File 2**).

### Gene silencing and CLIP-qPCR identify viral mRNA binders that promote viral infection

We silenced each of the 16 RBPs from the NP-mRNA core interactome with small interfering RNA (siRNA) pools in order to identify proteins with a pro- or antiviral activity. Non-target (NT) and NUP62-targeted siRNA pools were used as negative and positive controls, respectively (17). The siRNA-treated A549 cells were infected at a low MOI with a recombinant WSN-PB2-2A-Nanoluc influenza virus carrying a luciferase reporter gene (18), and luciferase activities were measured in cell lysates prepared at 24 hpi to monitor the efficiency of viral replication. A statistically significant 50 to 70% decrease of the luciferase signal compared to the NT control (p ≤ 0.001) was observed with five siRNA pools, targeted to members of the heterogeneous nuclear ribonucleoproteins family (hnRNPA2B1, hnRNPH1, TDP-43) or to splicing factors (SF3A3 and SF3B4) (**Figure 3A**). No strong reduction in cell viability was observed upon treatment of A549 cells with these siRNAs pools compared to the NT control (**Supp Figure 2A**). A 40% to 90% reduction of the luciferase signal was observed with at least three of the four siRNAs included in the initially tested siRNA pools (**Supp Figure 2B**), therefore making off-target effects very unlikely and indicating that hnRNPA2B1, hnRNPH1, TDP-43, SF3A3 and SF3B4 promote viral infection.

**Figure 3.**
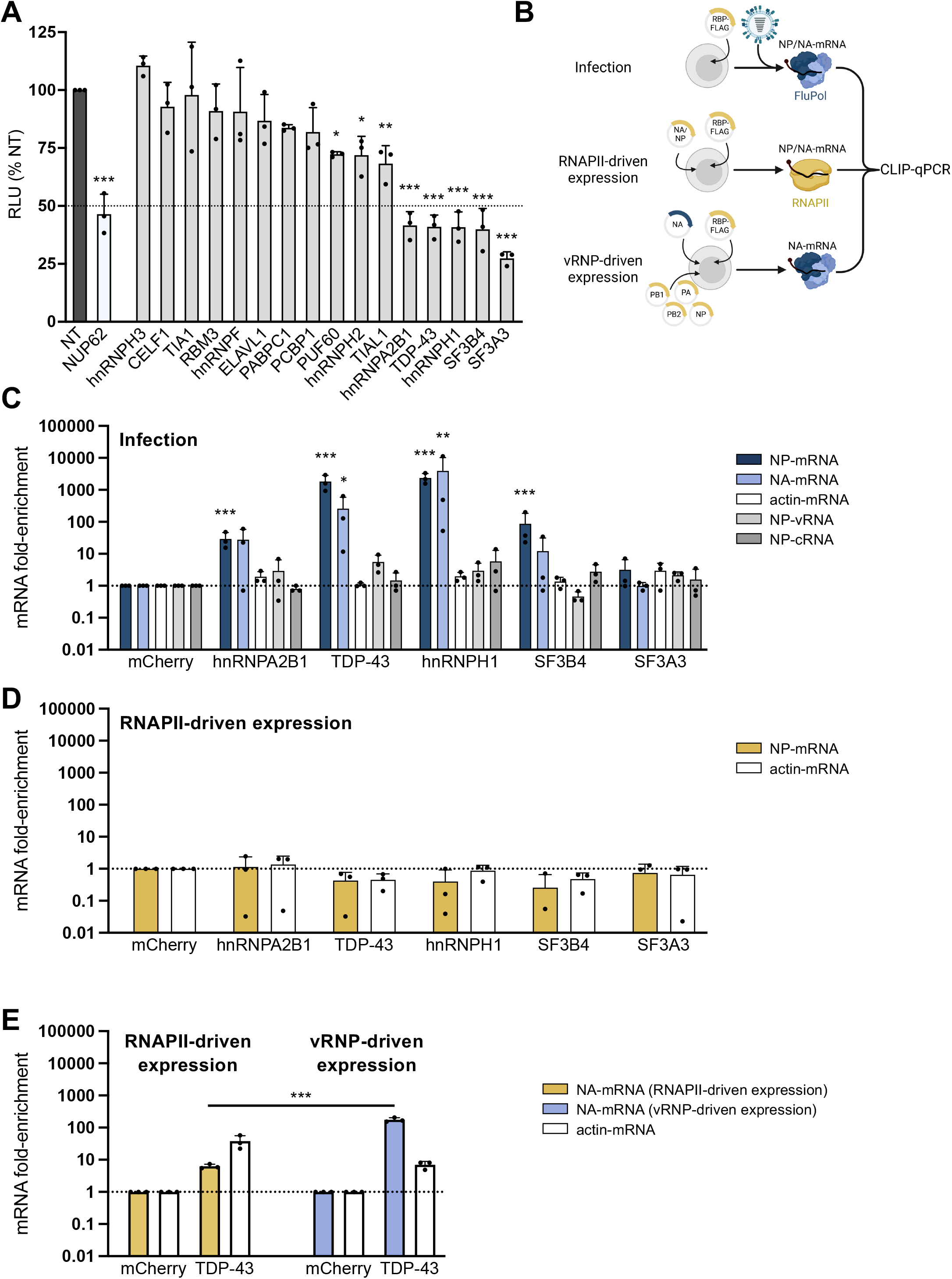
Functional analysis of the NP-mRNA core interactome. **A.** Secondary screening of the proteomics hits by siRNA-mediated silencing. A549 cells were treated with control non-target (NT, dark grey bar), NUP62 siRNAs (white bar) or siRNAs targeting the indicated RBPs (light grey bars) and infected with the WSN-PB2-2A-Nanoluc virus (0.001 PFU/cell). Luciferase activities were measured in cell lysates prepared at 24 hpi. Three independent experiments were performed in triplicates. The results (RLU: Relative Light Units) are expressed as the mean percentages ± SD of luciferase activity (100%: non-target siRNA). The significance was tested with a one-way ANOVA with Dunnett’s multiple comparisons test (reference: NT) using GraphPad Prism software (*p < 0.033; **p < 0.002; ***p < 0.001). **B.** Schematic representation of the CLIP-qPCR assays. HEK-293T cells were either i) transfected with the RPB-Flag expression plasmid and subsequently infected at a high MOI with the WSN virus (top); ii) co-transfected with the RPB-Flag expression plasmid and a pcDNA-NA or pcDNA-NP expression plasmid (middle); or iii) co-transfected with the RPB-Flag expression plasmid, pcDNA-PB1, PB2, PA and -NP plasmids, and a pPolI-NA plasmid that expresses the NA viral genomic RNA (bottom). In conditions i) and iii), viral mRNAs are transcribed by the viral polymerase (in blue), whereas in condition ii) they are transcribed by the cellular RNAPII (in ochre). At 6 hpi (i) or 24 hpt (ii and iii), cell lysates were prepared, and CLIP-qPCR was performed as described in the Methods section. **C-E.** Investigation of viral mRNA-binding proteins by CLIP-qPCR. (C) CLIP-qPCR was performed on HEK-293T cells overexpressing one of the candidate RBPs fused to the 3xFlag tag, or mCherry-3xFlag as a negative control. At 48 hpt HEK-293T cells were infected with WSN at a MOI of 5 PFU/cell for 6 h before UV crosslinking, lysis and pulldowns were performed. After the pulldown, RNAs cross-linked to RBPs were eluted from the beads by proteinase K digestion, followed by precipitation of the RNA, cDNA conversion and qPCR. The levels of NP-mRNA, -cRNA, -vRNA, NA-mRNA and actin-mRNA were determined, and their individual enrichments over the mCherry negative control were plotted. (D) To evaluate whether the interaction between any of the candidate RBPs and NP-mRNA was determined by the influenza transcription machinery CLIP-qPCR was performed on HEK-293T cells overexpressing individual Flag-tagged RBPs by transient transfection. Cells were co-transfected with an NP expression plasmid, where NP was transcribed from a RNPAII promoter. (E) To directly compare the association between a viral mRNA transcribed by the RNAPII or from a vRNP complex, HEK-293T cells were transfected with a mCherry-3xFlag or TDP-43-3xFlag expression plasmid, alongside either a RNAPII promoter-driven NA expression plasmid (ochre bars), or a PolI-driven NA vRNA expression plasmid together with expression plasmids for PB2, PB1, PA & NP, to reconstitute a vRNP (blue bars). At 48 hpt cells underwent CLIP-qPCR, and the levels of NA-mRNA and actin-mRNA were determined, and their individual enrichments over the mCherry negative control were plotted. The data shown are the mean ± SD of three independent experiments in triplicates. The significance was tested with a one-way ANOVA after log10 transformation of the data, with Dunnett’s multiple comparisons (reference: mCherry) (C-D) or with Tukey’s multiple comparisons test (E) using GraphPad Prism software (**p < 0.002; ***p < 0.001).

We performed Cross-Linking Immunoprecipitation-qPCR (CLIP-qPCR) on this subset of five RBPs. HEK-293T cells transiently expressing one RBP fused to a Flag tag were infected at a high MOI with the WSN virus (**Figure 3B**, top). The mCherry-Flag fusion protein was used as a negative control (**Supp Figure 3A**). As shown in **Figure 3C**, a significant 10- to 100-fold enrichment of co-immunoprecipitated NP-mRNA relative to the mCherry control was observed with hnRNPA2B1 and SF3B4, and a > 1000-fold enrichment was observed with TDP-43 and hnRNPH1 (dark blue bars), which underpins the robustness of our initial RNA interactome capture data. Notably, the same trends were observed for the viral NA-mRNA (**Figure 3C**, light blue bars). In the same conditions, no enrichment of the cellular actin mRNA, the viral genomic NP-vRNA or complementary NP-cRNA were observed (**Figure 3C**, white, light grey and dark grey bars respectively), suggesting a preferential recruitment of this RBP subset to viral mRNAs. CLIP-qPCR was repeated under conditions where synthesis of the viral NP mRNAs was under the control of the cellular RNA polymerase II (RNAPII) instead of the viral polymerase. To this end, HEK-293T cells were transfected with a RNAPII promoter-driven NP expression plasmid (pcDNA-NP), in addition to the indicated RBP-Flag or mCherry-Flag expression plasmid (**Figure 3B**, middle). Remarkably, no enrichment of NP-mRNA co-immunoprecipitating with the RBPs was observed in these conditions (**Figure 3D**, ochre bars), similar to what observed with the actin mRNA (**Figure 3D**, white bars), suggesting that the investigated RBPs are recruited to viral mRNAs through a mechanism which involves transcription by the viral polymerase.

### TDP-43 is recruited to viral mRNAs through interaction with the viral polymerase

We decided to focus on TDP-43, or TAR DNA-binding protein-43, which was initially identified as a repressor of HIV-1 transcription (56) and then found to be involved in the life cycle of several DNA and positive-stranded RNA viruses (57). It is a ubiquitous, 414 amino acids long RNA/DNA binding protein with a predominantly nuclear localization (58). Beyond its role as a transcription factor, TDP-43 regulates multiple aspects of mRNA metabolism, including splicing, nucleo-cytoplasmic transport, stability and translation (59). Mutations that cause misfolding and abnormal oligomerisation of TDP-43 are associated with an expanding spectrum of neurodegenerative diseases (60). There is also increasing evidence for a role of TDP-43 in human cancers (61).

To further investigate how TDP-43 is recruited to viral mRNAs, we performed CLIP-qPCR upon RNAPII-driven or vRNP-driven expression of viral mRNAs in a non-infectious setting. To this end, HEK-293T cells were transfected with a TDP-43-Flag expression plasmid in combination with a RNAPII promoter-driven NA expression plasmid (**Figure 3B**, middle), or with a mix of five plasmids allowing the reconstitution of vRNPs that actively transcribe viral NA-mRNAs (**Figure 3B**, bottom). Both conditions resulted in similar accumulation levels of the viral NA mRNA, as assessed by RT-qPCR in the cell lysates before immunoprecipitation (**Supp Figure 3B).** The enrichment of NA-mRNA co-immunoprecipitating with TDP-43 was significantly higher when the mRNA was synthesized by the vRNP-bound viral polymerase compared to when it was synthesized by the RNAPII (**Figure 3E**, blue and ochre bars, respectively), while the opposite trend was observed for the actin mRNA (**Figure 3E**, white bars).

Based on the CLIP-qPCR findings, we hypothesized that TDP-43 is recruited by the transcribing viral polymerase, and subsequently binds the viral mRNA to become assembled into mRNPs. We therefore investigated whether the viral polymerase and TDP-43 are physically interacting during the course of infection, using a split-luciferase based protein complementation assay either in an infectious setting (iPCA) (17), or alternatively in a transient pairwise expression setting (PCA) (**Figure 4A-B**). Two other viral mRNA binders validated by CLIP-qPCR, hnRNPH1, and SF3B4, were tested in parallel. Well-documented cellular partners of the viral polymerase, i.e. the RNAPII large subunit (RPB1) and the chicken ANP32A protein (chANP32A) were used as positive controls, whereas the TOP3B protein was used as a negative control based on a previous iPCA screen (17, 62, 63). Briefly, the RBPs of interest and control proteins, tagged with the Gaussia luciferase Gluc2 domain at their C-terminal end, were transiently expressed in HEK-293T cells. Transfected cells were then infected at a high MOI with a recombinant WSN influenza virus expressing a Gluc1-tagged polymerase (WSN-PB2-Gluc1) and luciferase activities were determined at 6 hpi. Normalized Luciferase activity Ratios (NLRs) with respect to background controls were determined as described in (17). The viral polymerase showed a positive interaction signal (≥ 3-fold the NLR observed with TOP3B) with TDP-43 as well as with hnRNPH1 and SF3B4 (**Figure 4A**). In a non-infectious context, i.e. when the viral polymerase and cellular RBPs were transiently co-expressed in the absence of other viral proteins or viral RNAs, an interaction could be observed only for TDP-43 and hnRNPH1, but not SF3B4, suggesting that other viral factors beyond the viral polymerase are needed for SF3B4 to associate with the viral polymerase (**Figure 4B**).

**Figure 4.**
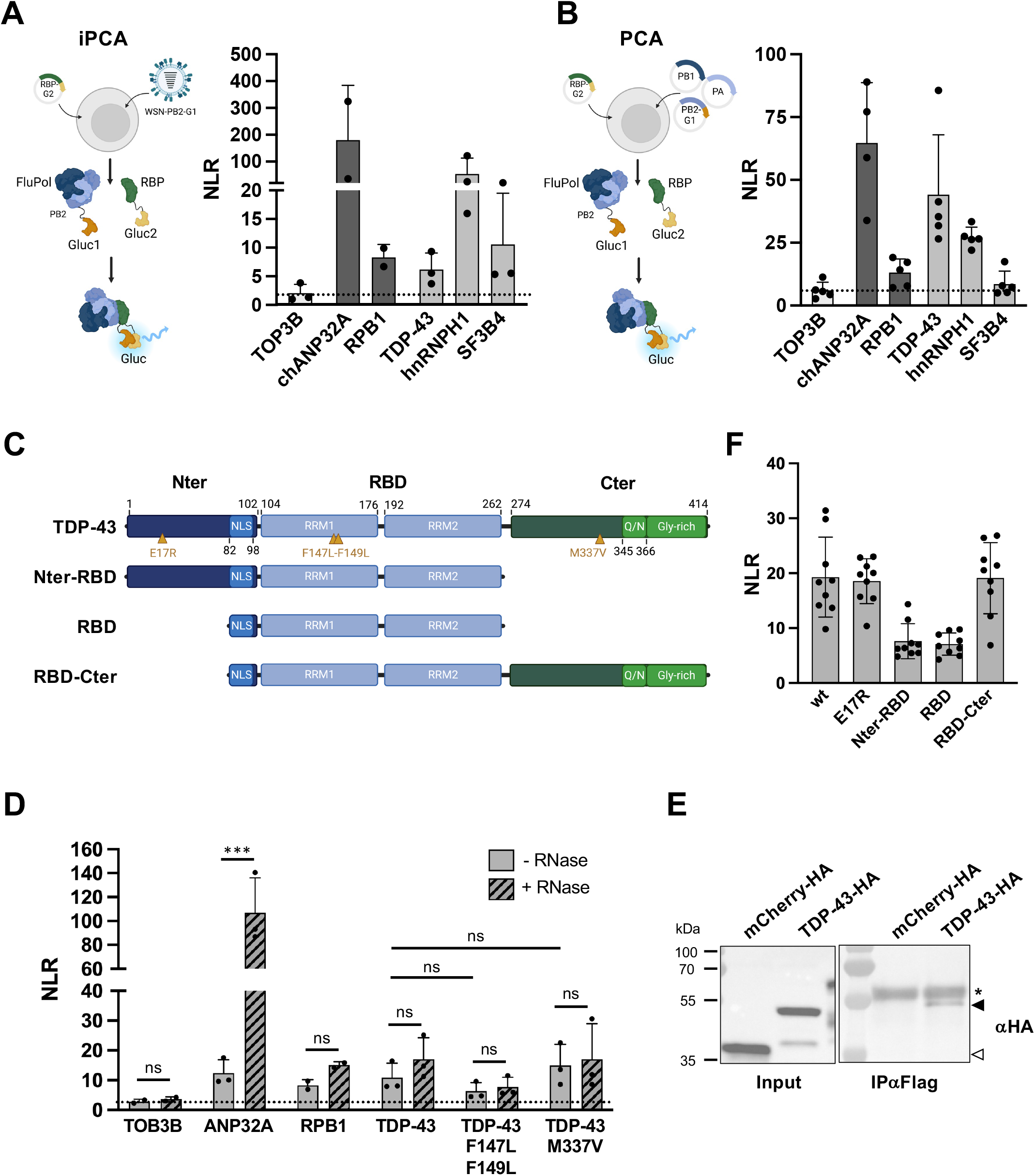
TDP-43 binds to the viral polymerase (FluPol). **A-B.** Cell-based RBP-FluPol binding assays. Schematic representation of the protein complementation assays based on two subdomains of the Gaussia luciferase (Gluc1/Gluc2 or G1/G2) are shown. HEK-293T cells were transfected with an expression plasmid for the proteins of interest fused to Gluc2 (RBP-Gluc2), and either infected with a recombinant WSN influenza virus expressing a Gluc1-tagged polymerase for iPCA (A) or co-transfected with expression plasmids for a WSN-derived, Gluc1-tagged viral polymerase for PCA (B). An active Gaussia luciferase is reconstituted when a direct RBP-FluPol interaction occurs. Luciferase activities were measured in iPCA cell lysates at 6 hpi (A) or in PCA cell lysates at 24 hpt (B). Normalised Luminescence Ratios (NLRs) were calculated as described in the Methods section. The data shown are the mean ± SD of three (A) or four (B) independent experiments performed in triplicates. The chicken ANP32A (chANP32A) and RPB1 controls were included in two of the three experiments shown in (A). The dotted line represents the NLR measured for the TOP3B protein, which was selected as a negative control based on previously published data (17). **C.** Schematic representation of TDP-43 subdomains and deletion mutants. Nter: N-terminal domain; RBD: RNA-binding domain; Cter: C-terminal domain; NLS: Nuclear Localization Domain; RRM1 and RRM2: RNA Recognition Motifs 1 and 2; Q/N: Glutamine/Asparagine-rich region; Gly-rich: Glycine-rich region. Amino-acid substitutions are indicated. **D.** RNase sensitivity of the interaction between TDP-43 and FluPol. Split-luciferase-based iPCA was performed as in (A), with the indicated cellular proteins fused to Gluc2. As an RNase-dependent interaction control, the Gluc2-tagged ANP32A and Gluc1-tagged viral NP proteins were co-expressed prior to infection with the wild-type WSN virus. Cell lysates were split in two halves which were supplemented with RNase A (solid bars) or not (hatched bars) and incubated for 30 min at 37°C prior to luciferase activity measurement. The data shown are the mean ± SD of three independent experiments performed in triplicates. The significance was tested with a two-way ANOVA with Sidak’s multiple comparisons test using GraphPad Prism software (***p < 0.001; ns: non significant). **E.** Co-immunoprecipitation of TDP-43 with the FluPol in the presence of RNase. HEK-293T cells were transfected for 24 h with HA-tagged constructs, then infected with the WSN-PA-3xFlag virus for 4 h. Cells were then harvested, lysed and subjected to Flag immunoprecipitation in the presence of RNase. A. Lysates (Input) and pull-down samples (IPαFlag) were analysed by western blot with an anti-HA antibody. Both the mCherry-HA control and TDP-43-HA are well expressed in the transfected cells, but only TDP-43-HA is detectable in the pull-down. The star denotes the heavy chain of the antibody used for the immunoprecipitation. Closed and open arrowhead : TDP-43-HA and mCherry-HA, respectively. **F.** Mapping of TDP-43 binding domain to the FluPol. Split-luciferase-based PCA was performed as in (B). The wild-type (wt) or indicated TDP-43 variants tagged with Gluc2 were transiently co-expressed with PB2-Gluc1, PB1 and PA expression plasmids derived from the WSN virus. The data shown are the mean ± SD of nine independent experiments performed in triplicates. The significance was tested with a one-way ANOVA with Dunnett’s multiple comparisons test (reference: wt or RBD-Cter) using GraphPad Prism software (* p< 0.033; **p < 0.002).

### The interaction of TDP-43 with the viral polymerase is RNA-independent and is mediated by its disordered C-terminal domain

TDP-43 binds RNA through two RNA recognition motifs, RRM1 and RRM2 (60), located in the central RNA binding domain (RBD) (**Figure 4C**). To investigate whether the interaction of TDP-43 with the viral polymerase is dependent on its RNA binding function, we tested the RRM1 mutant F147L-F149L, defective for RNA binding (64), along with the M337V mutant, reported to have a decreased affinity for G-quadruplex-containing mRNAs (65). When expressed transiently in HEK-293T cells, the mutants, fused to the V5-tag and the Gluc2 domain at their C-terminal extremity, showed similar steady-state levels as monitored by western-blot (**Supp Figure 4A**). Upon immunostaining with an anti-V5 antibody, the wild-type protein as well as the F147L-F149L and M337V mutants appeared predominantly nuclear, with a minor proportion of cells also showing a punctate cytoplasmic staining (**Supp Figure 4B**). When measured with the split-luciferase complementation assay, either in the infectious setting (**Figure 4D**, solid bars) or the transient expression setting (**Supp Figure 4C,** solid bars), the interaction signals measured between TDP-43 and the viral polymerase were not significantly affected by the presence of the F147L-F149L or M337V mutations, suggesting that TDP-43 RNA binding activity is not essential for the interaction with the viral polymerase. To investigate the potential need for bridging RNA molecules, the assay was performed in the presence or absence of exogenous RNase A. The co-expression of the human ANP32A (ANP32A) fused to Gluc2 and viral NP fused to Gluc1 was used as an RNase-sensitive interaction control (66). In the presence of RNase A, the interaction signal was increased in case of the ANP32-NP interaction whereas it was not significantly different for TDP-43-FluPol interactions, irrespective whether the wild-type or mutant TDP-43 proteins were expressed (**Figure 4D** and **Supp Figure 4C,** hatched compared to solid bars). Overall, our data show that the interaction of TDP-43 with the viral polymerase is RNA-independent.

The association between FluPol and TDP-43 was further assessed by co-immunoprecipitation, where HEK-293T cells were transfected with a plasmid to overexpress HA-tagged TDP-43 or mCherry 48 h prior to being infected at a high MOI with a recombinant influenza virus expressing a 3xFlag-tagged polymerase (WSN-PA-3xFlag). Cells were crosslinked at 4 hpi by PFA, then lysed in the presence of RNase A, and PA was immunoprecipitated using an anti-Flag antibody. Samples following immunoprecipitation, and corresponding input lysates, were analysed by Western blotting and the presence of HA-tagged TDP-43 or mCherry was determined (**Figure 4E**). TDP-43 was present in the immunoprecipitated samples while mCherry, which was expressed to a much greater level in the infected cells, was absent. Taken together, our observations show that the viral polymerase interacts with TDP-43 in influenza-virus infected cells and that this interaction is RNA-independent.

In addition to the central RBD domain, TDP-43 comprises an N-terminal homodimerization domain (NTD) and a disordered C-terminal domain (CTD). To map the FluPol binding domains of TDP-43, we tested the NTD-RBD, RBD and RBD-CTD deletion mutants (**Figure 4C**), as well as the E17R mutant, shown to be defective for TDP-43 dimerization (67). Importantly, the truncated proteins retained the nuclear localization domain (NLS, residues 82 to 98) located just upstream the RBD (68). Western-blot analysis showed that the variants accumulated at a slightly lower (RBD-CTD) or similar level compared to the wild-type (**Supp Figure 4A**), and their predominantly nuclear localisation was confirmed upon immunostaining (**Supp Figure 4B**). In the split luciferase-based assay, the N-ter-RBD and RBD mutants, but not the RBD-CTD or the E17R mutants, showed significantly decreased interaction signals compared to wild-type TDP-43 (**Figure 4F**). This shows that the interaction between TDP-43 and the viral polymerase does not involve the N-terminal domain of TDP-43, and suggests that the interaction is mediated primarily by the disordered C-terminal domain of TDP-43.

### Viral replication is impaired in TDP-43 knock-out cells

We achieved knock-out of TDP-43 in A549 cells by CRISPR/Cas9 as described in the Methods section. Briefly, lentiviral transduction was used first to establish a clonal cell line that stably expresses the Cas9 enzyme (referred to thereafter as Cas9 cells). In a second step, sgRNA targeting TDP-43 were introduced by lentiviral transduction (**Supp Figure 5A**). Two clonal cell lines expressing two distinct sgRNAs, referred to as TDP-43 KO1 and KO2, showed no detectable TDP-43 levels compared to the Cas9 cells as determined by western blot (**Figure 5A**, top lane). They were infected at a high MOI with the WSN virus and total cell lysates prepared at 4 and 6 hpi were analysed by western blot. The viral proteins (PB2, HA, NP and NS1) accumulated to lower levels in TDP-43 KO1 and KO2 cells compared to the Cas9 control (**Figure 5A**), consistent with our initial siRNA-depletion experiment (**Figure 3A**). Following infection of the TDP-43 KO2 and Cas9 cells at a low MOI with the wild-type WSN virus or with a representative human seasonal H1N1pdm09 influenza virus (A/Bretagne/7608/2009), plaque assays revealed a 1-log reduction in the yield of infectious viral particles in the supernatants of the KO2 compared to the Cas9 cells, at 24 hpi with the WSN virus and at 48 and 72 hpi with the H1N1pdm09 virus (**Figure 5B**). A similar trend was observed with a representative seasonal H3N2 virus (A/Centre/1003/2012) (**Supp Figure 5B**).

**Figure 5.**
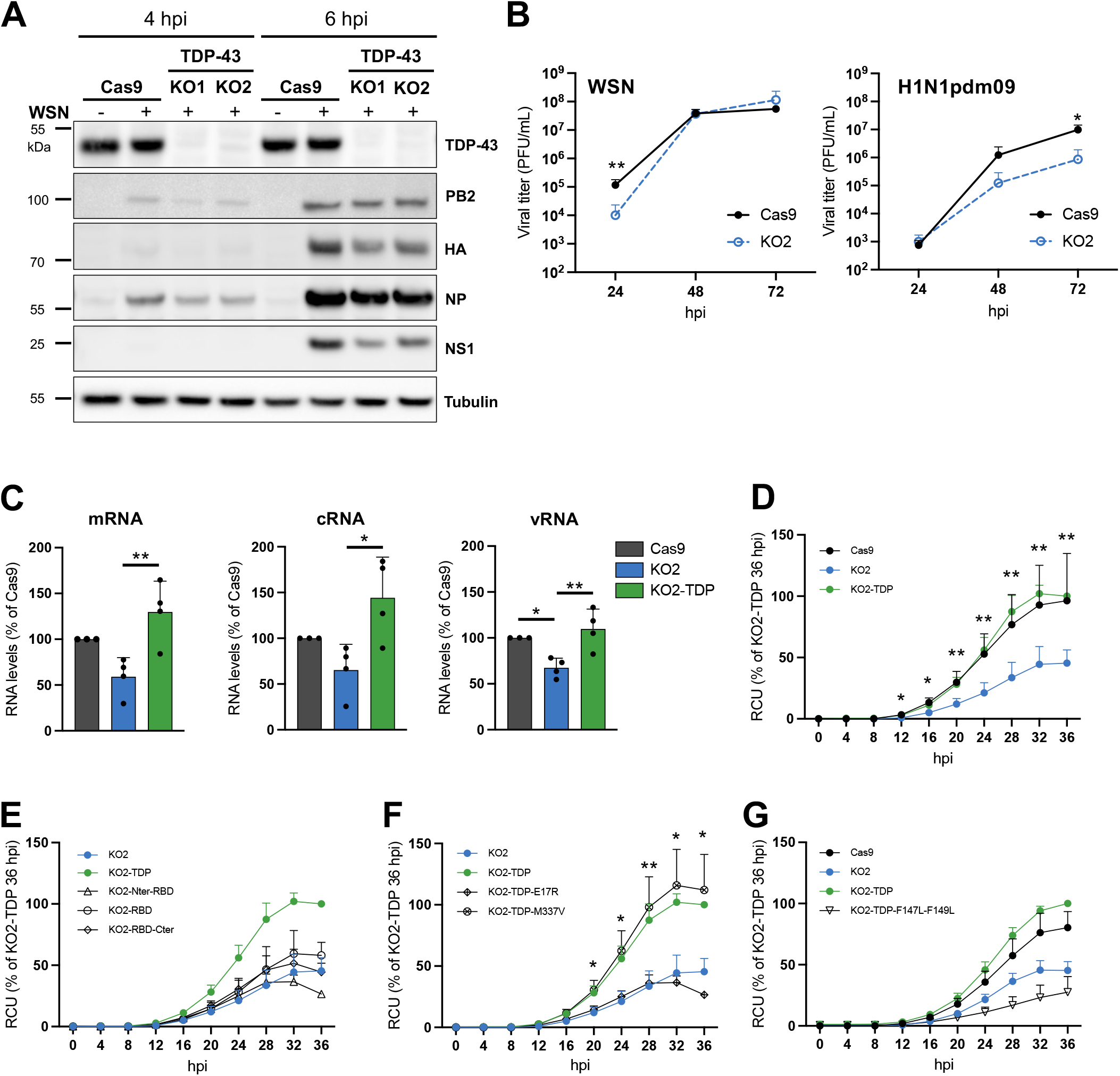
Viral growth is impaired in TDP-43 knock-out cells. **A.** Effect of TDP-43 knock-out on the steady-state levels of viral proteins. A549-derived control cells (Cas9) or TDP-43 knock-out cells (KO1 and KO2) were infected with the WSN virus at a MOI of 5 PFU/cell. Total extracts were prepared at 4 and 6 hpi and were analyzed by immunoblots using antibodies directed against the indicated proteins. Results representative of two independent experiments are shown. Cropped blots are shown. The corresponding full-length blots can be provided upon request. **B.** Effect of TDP-43 knock-out on the production of infectious viral particles. Cas9 or KO2 cells were infected with the WSN or a seasonal H1N1pdm09 virus at a low MOI. At 24, 48 and 72 hpi, the supernatants were collected and viral titers were determined by plaque assay. The data are expressed as the mean ± SD of three independent experiments in triplicates. Triplicate samples were pooled for titration. The significance was tested with a two-way ANOVA after log10 transformation of the data with Sidak’s multiple comparisons test using GraphPad Prism software (* p< 0.033; **p < 0.002). **C.** Effect of TDP-43 knock-out and rescue on the accumulation of viral NA RNAs. A549-Cas9, KO2 or KO2 cells rescued with the wild-type TDP-43 (KO2-TDP) were infected with the WSN virus at a MOI of 3 PFU/cell. Total RNAs were extracted at 5 hpi and subjected to strand-specific RT-qPCR as described in (42). RNA steady-state levels were normalised to GADPH and analysed using the 2^−ΔΔCT^ method (see Methods section) and are expressed as percentages (100%: Cas9 cells). The data shown are the mean ± SD of three independent experiments performed in triplicates. The significance was tested with a one-way ANOVA with Tukey’s multiple comparisons tests using GraphPad Prism software (* p< 0.033; **p < 0.002). **D-G.** Effect of gene rescue with wild-type or mutant TDP-43 proteins on viral growth. The control (Cas9, black curve, closed symbols), TDP-43 knock-out (KO2, blue curve) and wild-type (TDP, green curve) or mutant (NTD-RBD, RBD-CTD, RBD, E17R, M337V, F147L-F149L, black curves, open symbols) gene rescued cells were seeded on 96-well plates, infected with the WSN-PB2-2A-mCherry virus at a MOI of 10^-2^ PFU/cell and placed in the Incucyte S3 instrument for real-time monitoring of the fluorescence levels every 6 h up to 36 hpi. The data (RCU: Red Calibrated Units) are expressed as percentages (100%: KO2-TDP cells at 36 hpi) and are shown as the mean ± SD of three independent experiments performed in triplicates. The data in (D-F) were all generated in parallel, however for increased readability they are represented in three separate graphs, with the KO2 (blue curve) and wild-type gene rescue cells (green curve) being used as a common reference in all graphs. The significance (rescued versus KO2 cells) was tested with a two-way ANOVA with Dunnett’s multiple comparisons tests using GraphPad Prism software (*p < 0.033; **p < 0.002).

Wild-type TDP-43 or different mutants (F147L-F149L, E17R, M337V, RDB, NTD-RBD and NTD-CTD) were then re-expressed in the TDP-43 KO2 clone by lentiviral transduction followed by the selection of antibiotic-resistant polyclonal populations (**Supp Figure 5A**). Comparable steady-state protein levels were confirmed across the various transduced cell populations by western-blot, using an antibody specific for TDP-43 (**Supp Figure 5C** and **6A**, left panel) or the V5 C-terminal tag fused to TDP-43 (**Supp Figure 5C** and **6A**, right panel), except for the TDP-43 RBD mutant which accumulated at higher levels. Immunofluorescence staining with the anti-V5 antibody revealed a homogeneous proportion of stained cells with a predominantly nuclear staining except for the TDP-43 RBD and F147L-F149L mutants in which case an enhanced cytoplasmic localisation was observed (**Supp Figure 5D** and **6B**).

**Figure 6.**
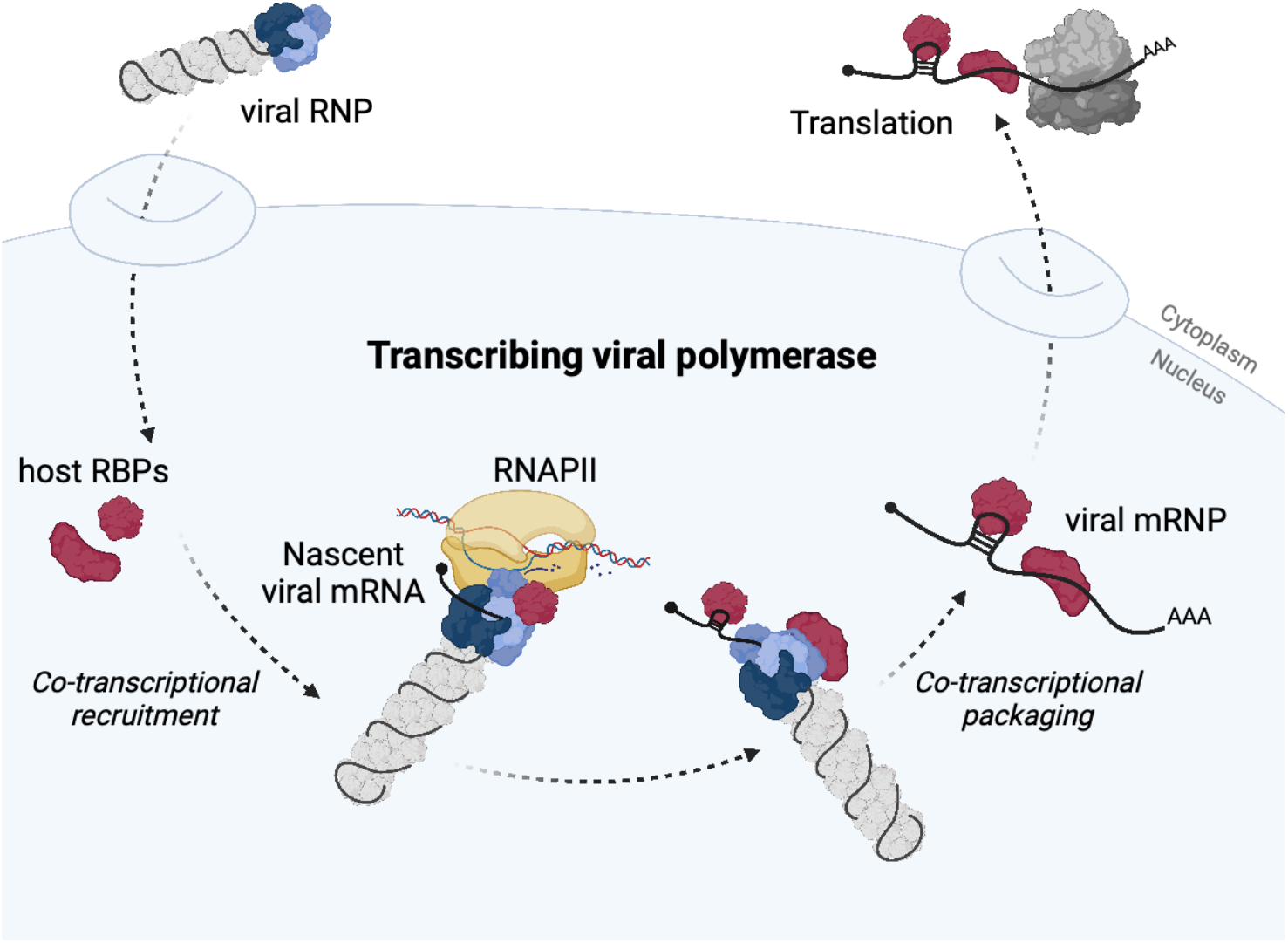
Model for the recruitment of host RNA binding proteins to viral mRNAs by the transcribing viral polymerase in the nucleus of influenza virus-infected cells. RNP: ribonucleoprotein. mRNP: messenger ribonucleoprotein. RBPs: RNA binding proteins. RNAPII: host RNA polymerase II.

We examined the effect of TDP-43 knock-out and rescue on the accumulation of NA viral RNAs upon infection at a high MOI, using a strand-specific RT-qPCR protocol (38). As shown in **Figure 5C**, the steady-state levels of m, c and vRNAs were reduced in TDP-43 KO2 cells compared to Cas9 cells and were restored upon wild-type gene rescue in the KO2-TDP cells, supporting the findings in western blot (**Figure 5A**) and plaque assays (**Figure 5B**). The rescued cell lines were then infected at a low MOI (10^-2^ PFU/cell) with a reporter WSN virus carrying a biscistronic PB2-2A-mCherry segment (69), and the mCherry fluorescence was monitored in real-time as described in the Methods section. The fluorescence levels were reduced on the TDP-43 KO2 compared to the Cas9 cells, and were restored in KO2-TDP cells (**Figure 5D**), in agreement with the data shown in **Figure 5C**. All investigated TDP-43 mutants (Nter-RBD, RBD, RBD-Cter, TDP-E17R, TDP-F147L-F149L) failed to rescue mCherry expression during WSN PB2-2A-mCherry infection as compared to the KO2 cells (**Figure 5E-G**, black curves), except for the M337V mutant (**Figure 5F**, crossed-circle symbols). Overall, the data demonstrate that TDP-43 specifically enhances influenza gene expression, and that all three domains of the protein, as well as homodimerization (TDP-E17R) and RNA-binding (TDP-F147L-F149L) are essential to its proviral action. Of note, we observed in contrast to previous reports that the knock-out of TDP-43 did not result in a higher level of cytoplasmic dsRNAs in KO2 compared to Cas9 and KO2-TDP cells (**Supp Figure 7A**), as assessed by immunofluorescence microscopy. Consistently, supernatants of poly(I:C)-treated Cas9, KO2 and KO2-TDP cells induced similar levels of Firefly luciferase activity when transferred to a reporter cell line which stably expresses the Firefly luciferase under the control of Interferon-Stimulated Response Elements (16) (**Supp Figure 7B)**, indicating that the Type I Interferon response is not significantly affected by TDP-43 depletion in A549 cells and is therefore not the underlying reason for the observed attenuation of viral growth.

## DISCUSSION

The RBPome of influenza virus mRNAs has so far been unexplored. We provide here the first characterisation of host proteins interacting with a specific influenza virus mRNA, namely NP-mRNA, during infection. Our workflow was designed to maximise the signal-to-noise ratio, by applying i) the low background eRIC method (14) with a single optimized LNA-modified capture probe; ii) differential analyses with respect to both mock-infected and non-crosslinked control samples in order to eliminate proteins bound non-covalently to beads or RNA (14); and iii) stringent statistical and fold-change cut-off values to define a core and an expanded interactome of 16 and 51 proteins, respectively. The high validation rate by CLIP-qPCR, with four out of five proteins of the core interactome being confirmed to bind viral positive-stranded mRNAs, but not viral negative- and positive-stranded genomic and antigenomic RNAs, supports the accuracy and specificity of our dataset.

Over twenty studies of viral RNA-host cell interactions on a proteome-wide scale have been published in the last decade, almost entirely conducted on single-stranded, positive-sense (ss+) RNA viruses, including several studies focused on SARS-CoV-2 (reviewed in (55)). The genomic vRNA of ss+ viruses also functions as a mRNA, as opposed to influenza and other negative-sense RNA viruses for which vRNAs and mRNAs are distinct, complementary species. About ∼200 proteins are present in the vRNA interactomes from three families of ss+ RNA viruses (*Coronaviridae*, *Togaviridae*, and *Flaviviridae*) (55), and 27 out of the 51 proteins from the influenza NP-mRNA interactome belong to this set of 200 proteins (**Supp File 2**). This observation suggests that specific mRNA processing pathways are hijacked by a very wide range of evolutionary distant viruses.

The NP-mRNA interactome reported here encompasses RBPs with known or potential roles in mRNA transcription (e.g. TDP-43, PUF60, TIAL1), splicing (e.g. SF3B4, TDP-43, hnRNPH1), transport (e.g. ELAVL1, hnRNPA1), stability (e.g. TDP-43, ELAVL1, hnRNPA2B1) and translation (e.g. GRSF1, PABP1, PCBP2). This is consistent with the fact that cross-linking was performed at 6 hpi, when NP-mRNAs are expected to exist in multiple functional stages including synthesis, nuclear export and translocation in the cytoplasm as well as translation. The NP-mRNA interactome shows little overlap with previously published genome-wide RNAi (70–74) and CRISPR (75, 76) screens aimed at identifying cellular factors involved in influenza virus life cycle. Still, PTBP1, RBM3, SNRPF, PTMA, RPS6, hnRNPA1, hnRNPDL and NUDT21 were previously reported as primary hits in at least one these genome-wide functional screen. Our own focused RNAi screen targeting the 16 components of the NP-mRNA core interactome revealed five proteins with proviral activity (hnRNPA2B1, hnRNPH1, TDP-43, SF3B4 and SF3A3). The relatively low number of proteins identified as having pro- or antiviral function might be due to functional redundancy among the RBPs and/or to the fact that the knock-down was not effective enough in some instances.

Five proteins from the expanded NP-mRNA interactome, GRSF1, hnRNPK, hnRNPA2B1, NUDT21 and PABPC1 were previously documented as being involved in the IAV life cycle. GRSF1 and PABPC1 were found to bind the 5’ UTR of viral mRNAs including the NP-mRNA and to stimulate their translation (44, 45, 51, 54, 77), further supporting the robustness of our dataset. The hnRNPK protein was previously shown to bind the viral M1-mRNA and to regulate its splicing jointly with the NS1-BP protein (46, 47). Our data suggest that hnRNPK also regulates the fate of unspliced viral mRNAs such as the NP- and NA-mRNAs. The NUDT21 protein was found to interact with the influenza PA-X protein and to contribute to its RNase activity towards host RNAs (50). The depletion of hnRNPA2B1 was reported in two independent studies to induce a two-to ten-fold increase in the production of infectious viral particles (48, 49), which contrasts with our finding of a decreased viral replication. This discrepancy is most likely related to differences in the cell lines, virus strains and/or experimental read-outs.

Although influenza mRNAs are capped and poly-adenylated by unique mechanisms, all except one of the 51 proteins bound to the influenza NP-mRNA have already been identified as being part of the global interactome of human poly(A) mRNAs (9, 10). Interestingly, the viral RPBome represents a strongly interconnected network of protein according to the STRING database. Indeed, each protein interacts on average with ∼40% of other proteins within the viral RBPome, with hnRNP proteins showing the highest degree of connectivity. In contrast, each protein interacts on average with only ∼4% of other proteins within the group of 658 RPBs found to bind cellular mRNAs in two independent studies (9, 10), with ribosomal proteins being mostly involved (**Supp File 2**). The 51 host proteins interacting with influenza virus NP-mRNA could possibly be representative of a particular subset of cellular mRNPs that share a similar protein composition and function. Indeed, some evidence suggest that mRNA processing is a highly controlled, RNP-driven process (78). However, the biochemical characterisation of individual endogenous cellular mRNPs using in *vivo* RNA-centric approaches remains challenging and very few datasets are available (11). To our knowledge, the only human endogenous mRNP characterised so far is the c-myc mRNP (79). Only five proteins of the influenza NP-mRNA interactome (hnRNPAB, PABPC1, ELAVL1, SRSF2, RPS6) are found among the list of 229 proteins reported to bind the c-myc mRNA. Conversely, none of the hnRNPA2B1, hnRNPH1, TDP-43, SF3B4 or SF3A3 NP-mRNA binders was found to bind the cellular actin mRNA in the present study. Much broader comparisons between the interactomes of individual mRNAs will be needed in the future to uncover common and distinct regulatory mechanisms among cellular and viral mRNAs.

To our knowledge, so far only one study has investigated viral RNA-host protein interactions in influenza virus-infected cells on a proteome-wide scale, using the viral cross-linking and solid-phase purification or VIR-CLASP approach (80). Influenza virus particles containing 4SU-labelled vRNAs were used for high MOI infection followed by UV-crosslinking at a very early time point, to allow the pull-down and identification of cellular proteins that associate with the incoming vRNAs. Intriguingly, the ∼700 proteins reported to bind the incoming genomic vRNAs comprise 14 among the 51 proteins we identified in the NP-mRNA interactome (**Supp File 2**). These 14 proteins include the TDP-43 and hnRNPA2B1 proteins, for which our CLIP-qPCR experiments confirmed the binding to viral mRNAs but did not reveal any significant binding to genomic vRNAs in influenza virus infected cells at 6 hpi. This discrepancy could possibly be due to methodological differences (e.g. 4SU-labelled RNAs and a MOI of 10^3^ PFU/cell in the VIR-CLASP study, versus unlabelled RNAs and a MOI of 10 PFU/cell in our study). Alternatively, it could reflect temporal differences in the interactomes of influenza genomic vRNAs, depending on whether cross-linking was performed at an early or a later time point after infection. A limitation of our study as well as most studies published so far is that they are limited to a single time-point, and do not separate nuclear from cytoplasmic RNAs. Synchronized infections, cross-linking at various time points or inhibition of viral replication at specific stages of infection, in conjunction with cell fractionation, will be needed to document precisely the temporal dynamics of viral RNA interactomes.

Unexpectedly, we demonstrate that the viral polymerase plays an essential role in the assembly of viral mRNPs by recruiting RBPs through direct or close interactions. Indeed, we show that viral mRNA are bound by TDP-43, hnRNPA2B1, hnRNPH1 and SF3B4 when synthesized by the viral polymerase in infected cells, but not when synthesized by the cellular RNAPII transcription apparatus. Likewise, viral mRNAs are bound by TDP-43 when synthesized by a transiently reconstituted vRNP, i.e. in the absence of any other viral proteins than the viral polymerase and NP. The fact that TDP-43 was found to find the viral NP-mRNA as efficiently as the viral NA-mRNA, and the same was observed for hnRNPA2B1, hnRNPH1 and SF3B4, supports the model that these proteins are recruited primarily through protein-protein interactions with the viral polymerase before binding the mRNAs, possibly through primary sequence motifs and/or secondary structures. There is multiple *in vitro* and *in vivo* evidence that TDP-43 specifically binds UG-rich RNA sequences in the 3’ UTR or the introns of various genes (81–84). The GUGUG and UGUGU motifs, found to be strongly enriched among TDP-43 binding RNA sequences upon iCLIP experiments (82), are present on all full-length mRNAs of the influenza WSN strain at one to eight copies per mRNA. Whether some of these motifs are actually bound by TDP-43 warrants further investigations.

Our data reveal that the TDP-43 protein promotes influenza virus replication. Within our proteomics dataset, TDP-43 was among the strongest hits as it showed a ∼2.5- and 4-fold enrichment in infected versus mock and in crosslinked versus non-crosslinked samples, respectively. It also showed strong binding to the viral NP- and NA-mRNAs in subsequent CLIP-qPCR experiments, with an observed ∼1000 fold-enrichment compared to the mCherry control. CRISPR-mediated KO of TDP-43 in A549 cells resulted in a moderate but reproducible reduction in the accumulation of viral RNAs and proteins, which was reversed upon TDP-43 rescue. Moreover, we demonstrate that the interaction between TDP-43 and the viral polymerase is RNA-independent, does not involve the N-terminal domain of TDP-43 and could be mediated primarily by its disordered C-terminal domain.

Taken together, we propose a model in which the viral polymerase recruits TDP-43 to the viral mRNAs (**Figure 6**). In agreement with previously published RNA co-immunoprecipitation assays (54), our mass-spectrometry findings confirm that the viral polymerase itself is not part of the viral mRNPs. However, our data strongly supports the hypothesis that the viral polymerase binds TDP-43 through an RNA-independent protein-protein interaction and mediates its co-transcriptional recruitment on nascent viral mRNAs. The physical association of the viral polymerase with the cellular RNAPII for cap-snatching could possibly facilitate the recruitment of TDP-43. Indeed, beyond its well-established role in post-transcriptional RNA processing, TDP-43 also appears to play a role in regulating the transcription of human genes (85–87). The model we propose for TDP-43 recruitment to viral mRNAs may be applicable to hnRNPH1, hnRNPA2B1, and SF3B4, which would confer a central role to the viral polymerase for the recruitment of mRNA processing factors. A common feature of these four proteins is that they have long C-terminal disordered domains, known to serve as scaffolds for dynamic protein-protein interactions and the formation of biomolecular condensates, and to be a common feature of RNAPII-related transcription factors (88, 89). Our finding that these proteins get assembled into influenza mRNPs might serve as a starting point to explore to what extent phase-separation plays a role in transcription and post-transcriptional processing of viral mRNAs.

## ACCESSION NUMBERS

The mass spectrometry proteomics data have been deposited to the ProteomeXchange Consortium via the PRIDE (25) partner repository with the dataset identifier PXD040783.

## SUPPLEMENTARY DATA STATEMENT

Supplementary Data will be made available at NAR online. They are submitted as a separate file.

## ACKNOWLEDGEMENTS

We would like to thank Dr. Feng Zhang (Broad Institute, Cambridge, USA), Dr. Mikhael Matrosovich (Philipps Universität, Marburg, Germany), Dr. Pierre-Olivier Vidalain (CIRI, Lyon), Dr. Daniel Marc (INRAE, Nouzilly, France), Dr. Caroline Demeret, Dr. Sandie Munier and the National Reference Center for Respiratory Viruses (all at Institut Pasteur, Paris, France) and for sharing reagents used in this work. We would like to thank Yves Jacob, Anastasia Komarova and Valérie Najburg (Institut Pasteur, Paris, France) for helpful discussions. Some figures were created with BioRender.com.

