## Supplemental Figures 1-7 for "The RBPome of influenza A virus mRNA reveals a role for TDP-43 in viral replication"

### Supp Figure 1

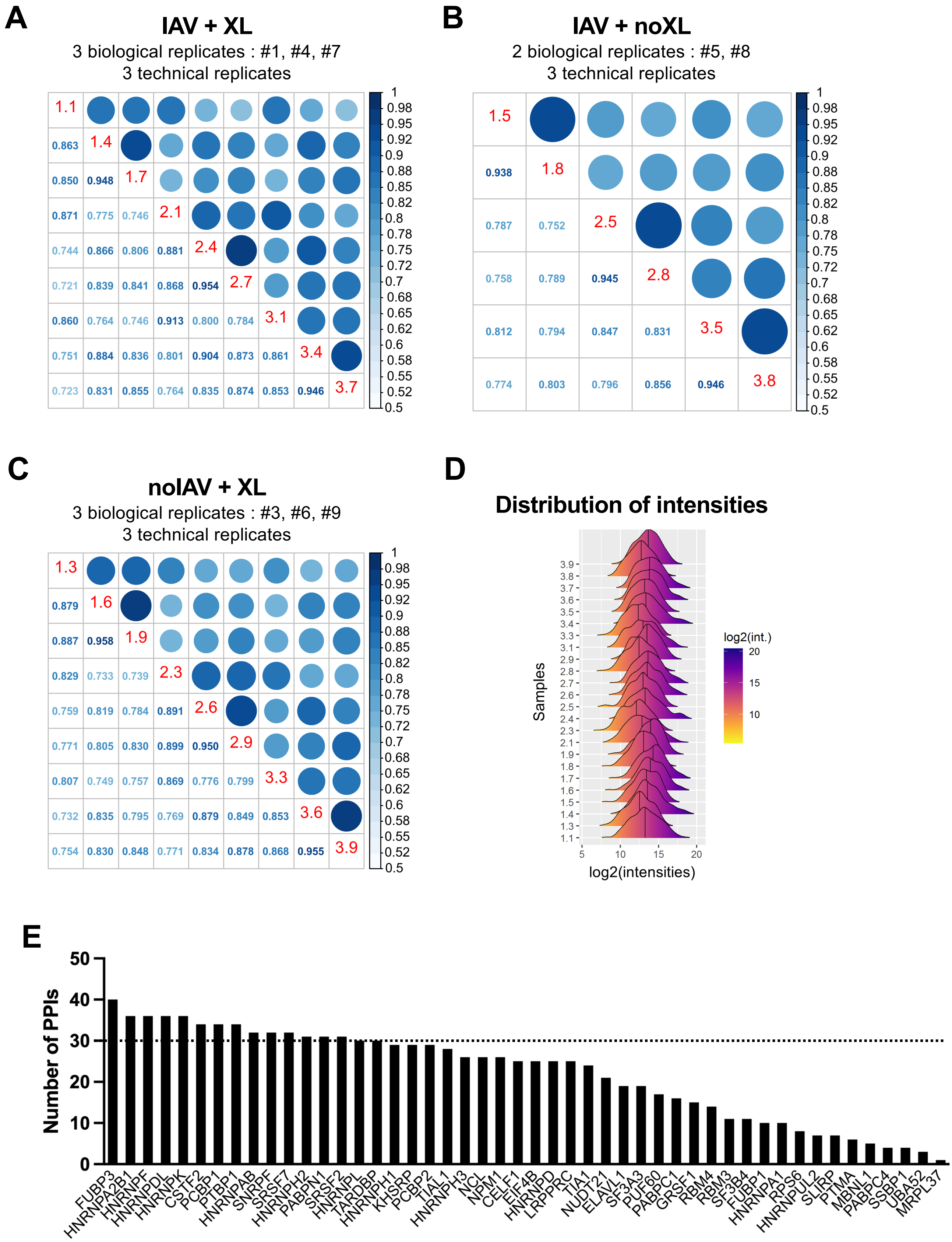

**Supp Figure 1. Identification of cellular proteins bound to influenza virus NP-mRNA in human infected cells (related to Figures 1 and 2).**

**A-C.** Correlation matrices between replicates of the different conditions. A correlation matrix represent the Pearson correlation coefficients between each pair of samples computed using all complete pairs of intensity values measured in these samples. Intensity values correspond to TMT-MS2 quantitative relative abundance metrics in the columns titled “Reporter intensity corrected” of the "proteinGroups.txt" file of MaxQuant. The samples identification numbers are indicated in red on the diagonals in the format “number of the technical replicate.number of the biological replicate”. Pearson correlation coefficients are indicated in the lower triangular parts of the matrices. In the upper triangular parts, the diameters and gradient colors of the circles are function of these coefficients.

**D.** Distributions of the log2(intensities) of proteins in all proteomics samples. Intensity values correspond to TMT-MS2 quantitative relative abundance metrics in the columns titled “Reporter intensity corrected” of the "proteinGroups.txt" file of MaxQuant. The samples identification numbers are indicated on the vertical axis in the format “number of the technical replicate.number of the biological replicate”.

**E.** Degree of connectivity among proteins of the expanded interactome. For each protein, the number of predicted protein-protein interactions within the expanded interactome is indicated.

### Supp Figure 2

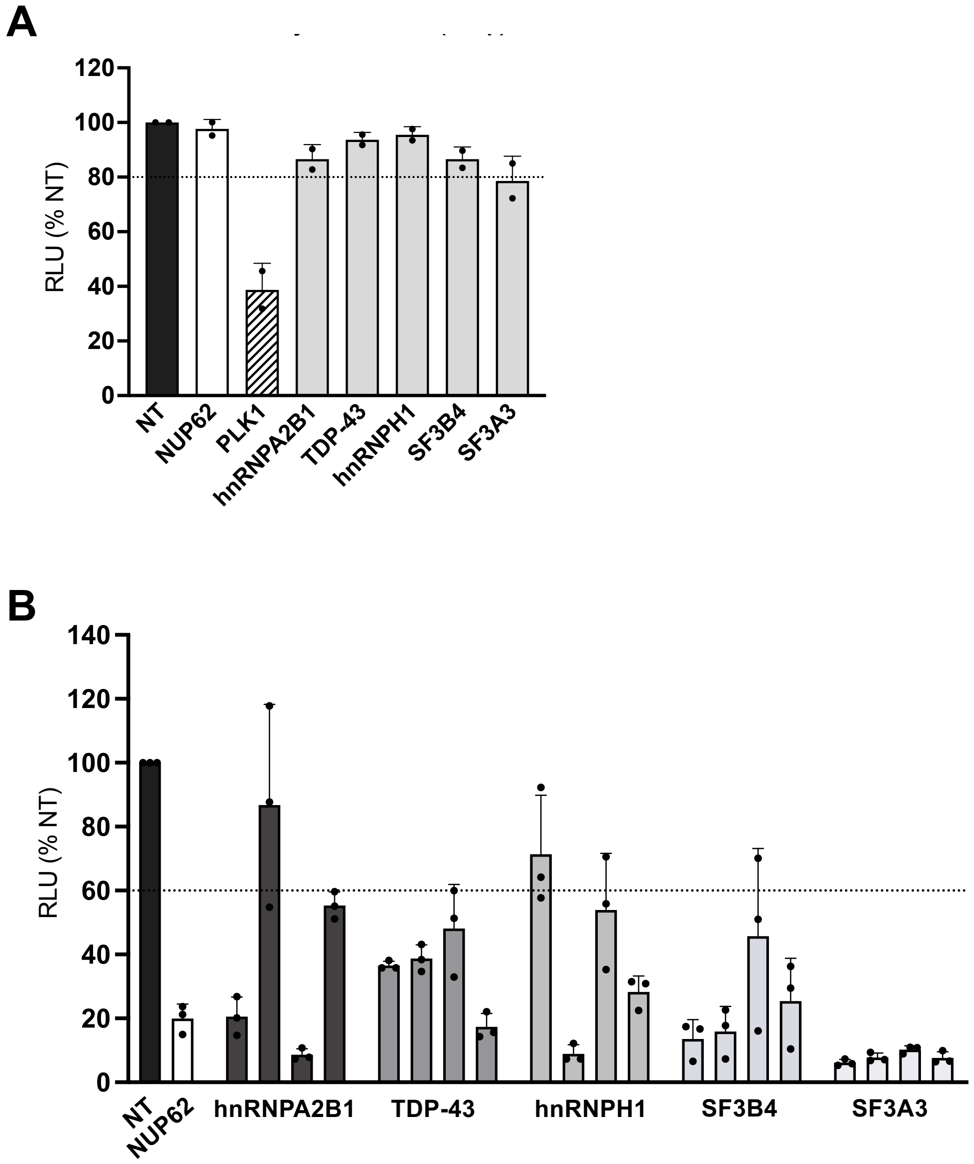


**Supp Figure 2. Gene silencing of RPBs (related to Figure 3A).**

**A.** Toxicity of siRNA pools. A549 cells were transfected with 37.5 nM of siRNA and cell viability was determined at 72 hpt using the CellTiter-Glo Luminescent Viability Assay kit (Promega). siRNAs targeting PLK1 kinase was used as a positive control (hatched bar). The data shown (RLU: Relative Light Units) are the mean ± SD of two independent experiments performed in triplicates and are expressed as percentages (100%: NT siRNA). The dotted line indicates a 20% reduction in luciferase signal, considered as a threshold for significant toxicity.

**B.** Exclusion of off-target RNAi effects by using deconvoluted siRNA pools. A549 cells were treated with control non-target (NT, black bar) or NUP62 siRNAs (white bars) or with each of the four individual siRNAs that constitute the siRNA pools tested in Figure 3A (grey bars). After 48 h transfected cells were infected with the WSN-PB2-2A-Nanoluc virus (0.001 PFU/cell). Luciferase activities were measured in cell lysates prepared at 24 hpi. The data shown are the mean ± SD of three independent experiments performed in triplicates and are expressed as percentages (100%: NT siRNA). The dotted line indicates a 40% reduction in luciferase activity.

### Supp Figure 3

**A**

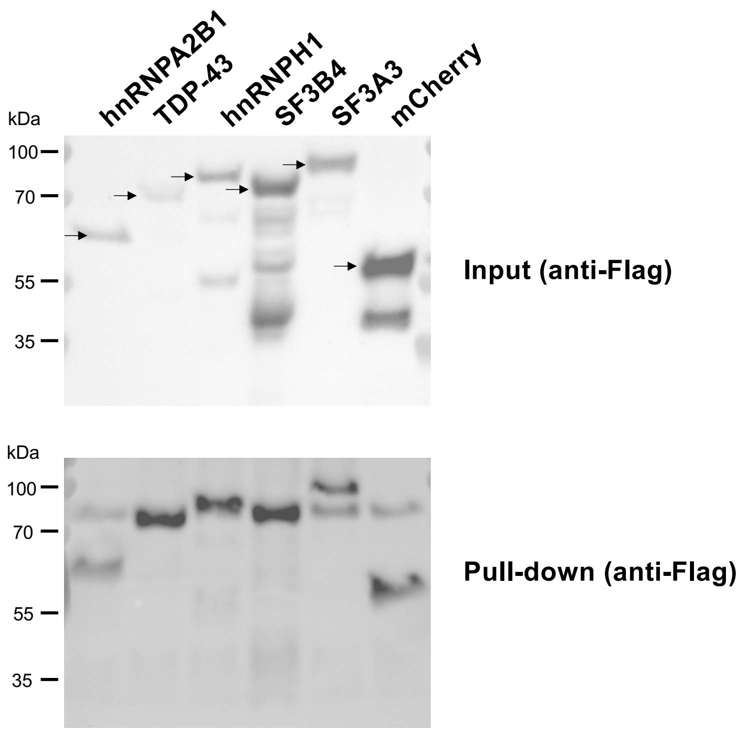


**B**

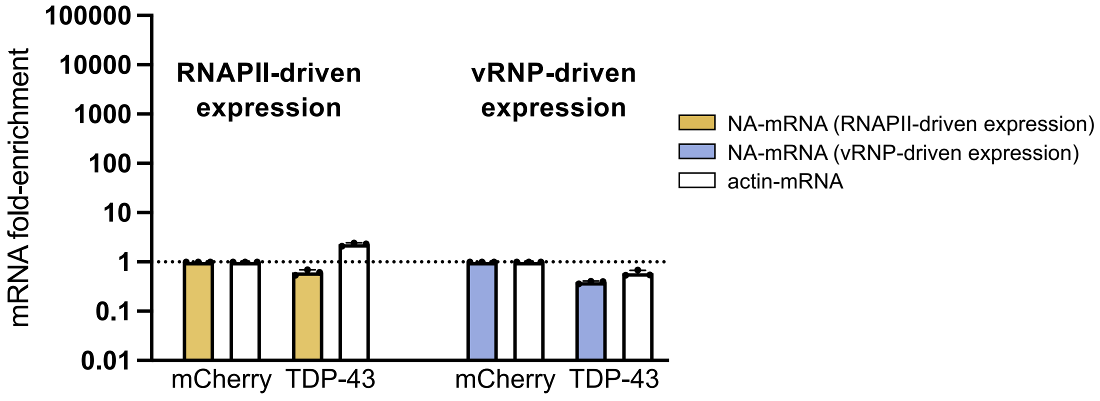


**Supp Figure 3. Investigation of viral mRNA-binding proteins by CLIP-qPCR (related to Figure 3B-E)**

**A**. Western blot showing the level of each individual 3xFlag-tagged RBP at the time of harvesting for CLIP-qPCR. Upper panel shows the input, while the lower panel shows the proteins that were pulled down. Arrows indicate the band corresponding to the correct molecular weight for each RBP.

**B.** qPCR of the input RNA levels for the pulldowns performed in Figure 3E showing comparable levels of NA- and actin-mRNA between the RNAPII and the vRNP-driven expression conditions. The data shown are the mean ± SD of three independent experiments in triplicates.

### Supp Figure 4

A

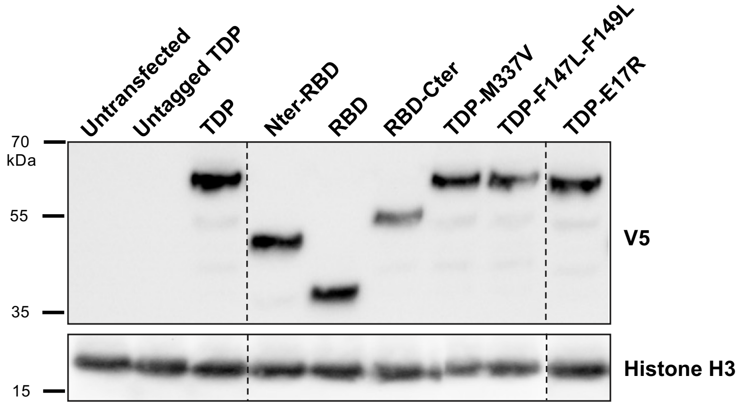


B

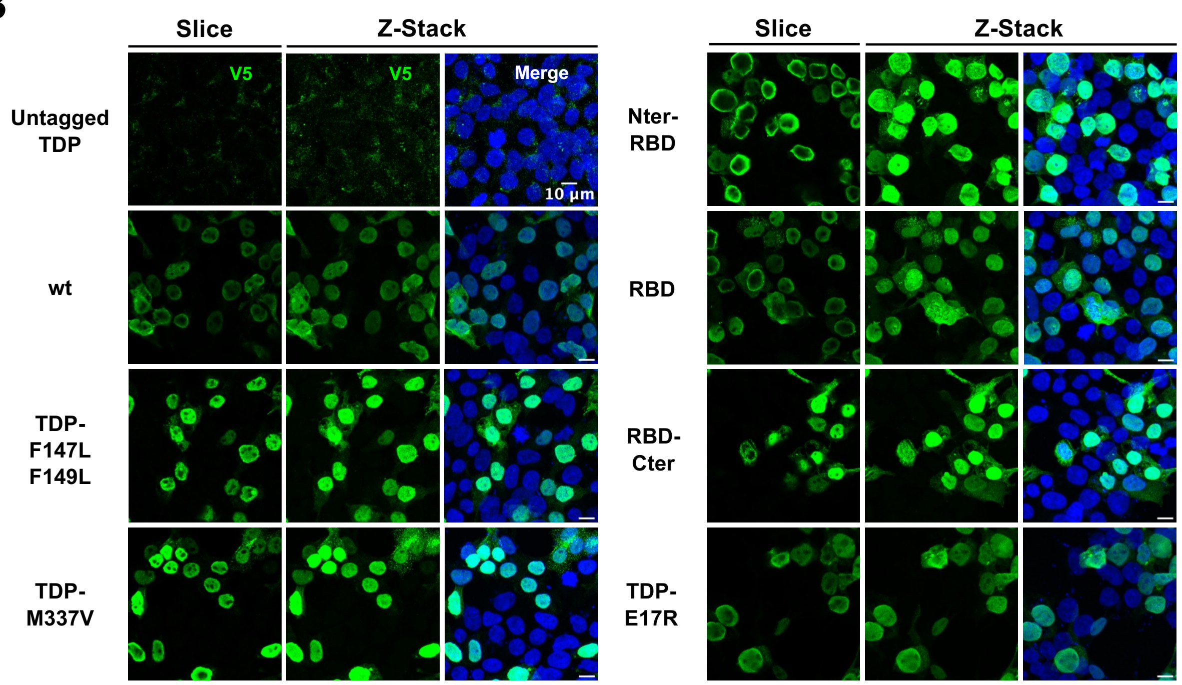


C

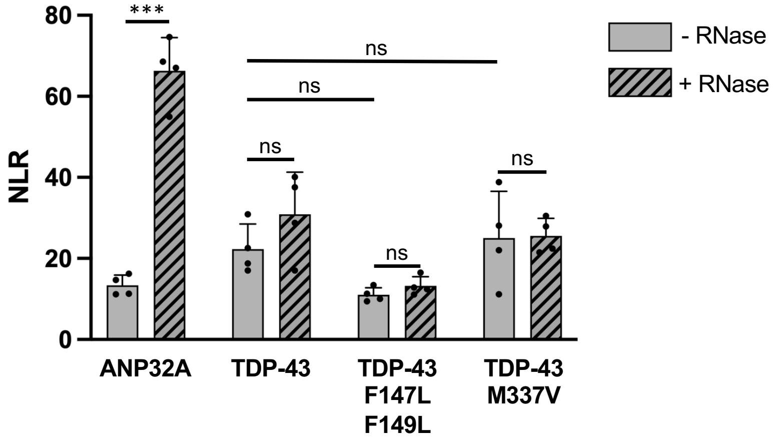

**Supp Figure 4. Characterisation of TDP-43 variants (related to Figure 4).**

**A.** Steady state levels of the recombinant wild-type and variant TDP-43 proteins upon transient expression in HEK-293T cells. HEK-293T cells were transfected with the expression plasmids for the indicated proteins, fused to a V5 tag at their C-terminal end. Total extracts were prepared at 24 hpt and were analyzed by immunoblot using an antibody directed against the V5 tag. The membranes were re-hybridized with an anti-Histone H3 antibody to control for the total protein load. The dashed lines indicate the juxtaposition of non-adjacent lanes from the same immunoblot.

**B.** Immunostaining of the recombinant wild-type and TDP-43 proteins upon transient expression in HEK-293T cells. HEK-293T cells were seeded on coverslips and transfected with the indicated expression plasmids. Cells were fixed at 24 hpi and were stained with an antibody against the V5-tag to visualise the V5-tagged recombinant TDP-43 proteins and with DAPI to visualise the nuclei. Representative images are shown. Z-Stack: stacking of 20 consecutive optical slices 0.35 µm thick. Slice: a single optical slice. Scale bar: 10 µm.

**C.** RNase sensitivity of the TDP-43-FluPol interaction. Split-luciferase-based PCA was performed as in Figure 4C, with a combination of the indicated wild-type and mutant TDP-43 cellular proteins fused to Gluc2 and a Gluc1-tagged FluPol, or a combination of Gluc2-tagged ANP32A and Gluc1-tagged NP. Prior to luciferase activity measurement, the cell lysates were split in two halves which were supplemented with RNase A (solid bars) or not (hatched bars), and incubated for 30 min at 37°C. The data shown are the mean ± SD of three independent experiments performed in triplicates. The significance was tested with a two-way ANOVA with Sidak's multiple comparisons test using GraphPad Prism software (***p < 0.001; ns: non significant).

### Supp Figure 5

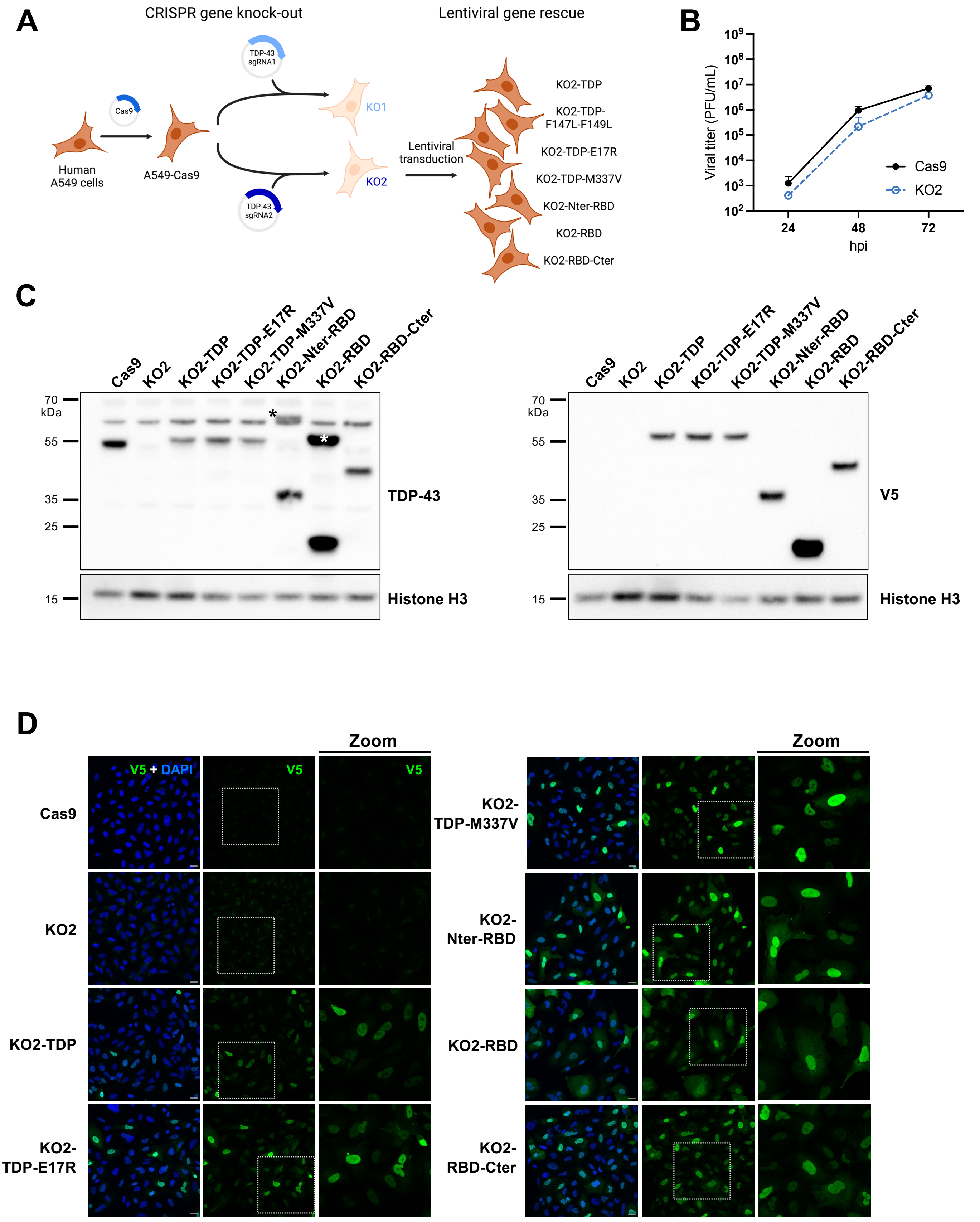

**Supp Figure 5. TDP-43 silencing and rescue (related to Figure 4).**

**A.** Schematic representation of TDP-43 CRISPR knock-out and lentiviral rescue.

**B.** Effect of TDP-43 knock-out on the production of infectious viral particles. A549-Cas9 (Cas9) or KO2 cells were infected with a seasonal H3N2 virus at a low MOI. At 24, 48 and 72 hpi, the supernatants were collected and viral titers were determined by plaque assay. The data are expressed as the mean ± SD of two independent experiments. Triplicate samples were pooled for titration. Differences were found not significant when tested with a two-way ANOVA after log10 transformation of the data using GraphPad Prism software.

**C.** Steady state levels of the recombinant wild-type and mutant TDP-43 proteins upon lentiviral rescue of the KO2 cells. Total extracts were prepared from ~ 15 x $10^4$ cells and were analysed by immunoblot using an antibody directed against TDP-43 (left panel) or the V5-tag (right panel). Membranes were re-hybridized with an anti-Histone H3 antibody to control for the total protein load. Unexplained high molecular weight bands revealed with the anti-TDP-43 antibody in the KO2-TDP-Nter-RBD and KO2-TDP-RBD samples are indicated by a black and a white star, respectively.

**D.** Immunostaining of the recombinant wild-type and mutant TDP-43 proteins upon lentiviral rescue of the KO2 cells. The indicated cell lines were seeded on glass coverslips and stained with an antibody against the V5-tag to visualise the V5-tagged recombinant TDP-43 proteins and with DAPI to visualise the nuclei. Representative images are shown. Z-stack images are covering 14 consecutive optical slices of 0.35 µm thick. Scale bar: 20 µm.

### Supp Figure 6

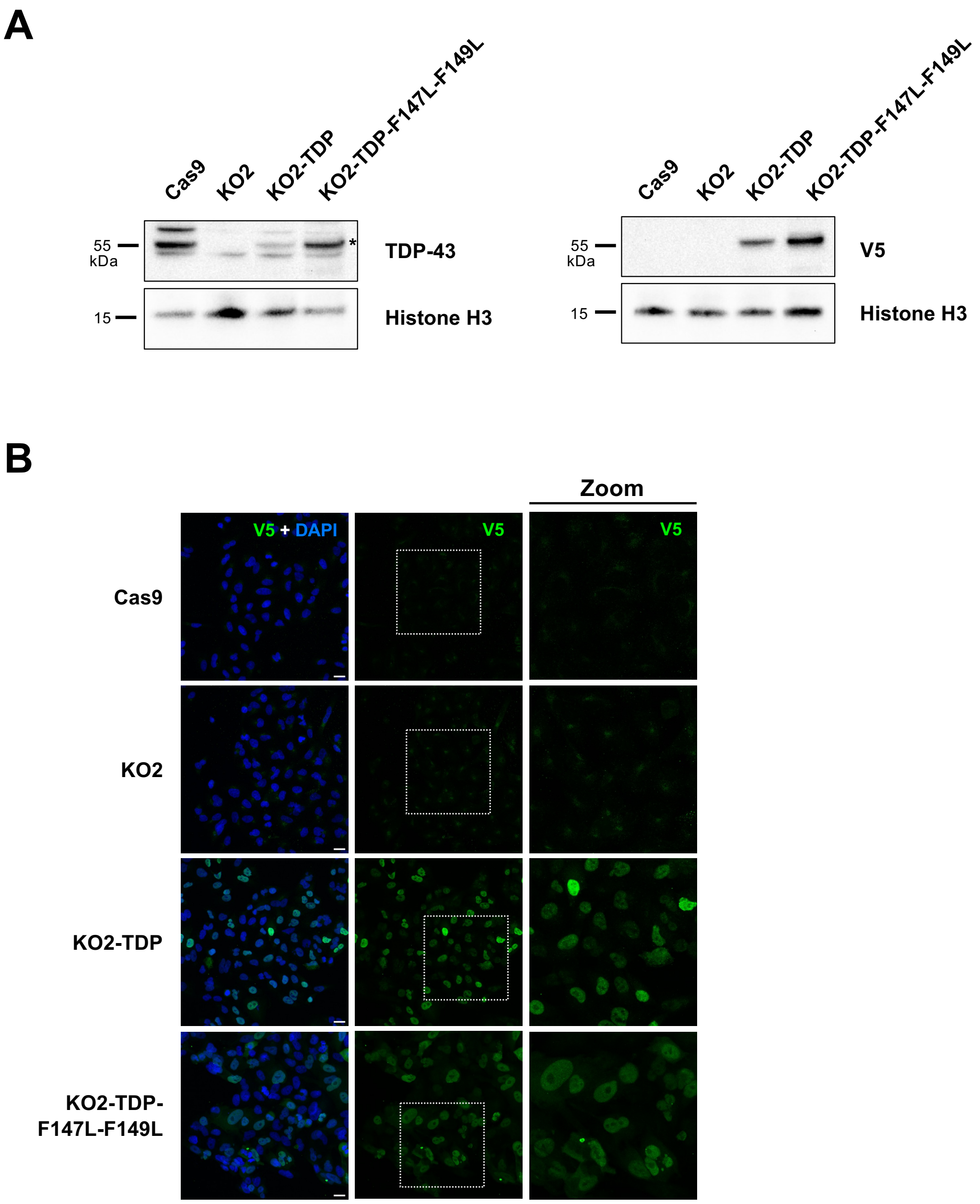


**Supp Figure 6. TDP-43 silencing and rescue with the TDP-43-F147L-F149L mutant (related to Figure 4).**

**A.** Steady state levels of the recombinant wild-type and mutant proteins upon lentiviral rescue of the KO2 cells. Total extracts were prepared from ~ 15 x $10^4$ cells and were analysed by immunoblot using an antibody directed against TDP-43 (left panel) or the V5-tag (right panel). The membranes were re-hybridized with an anti-Histone H3 antibody to control for the total protein load. The band corresponding to TDP-43 is indicated by a star.

**B.** Immunostaining of the recombinant wild-type and mutant proteins upon lentiviral rescue of the KO2 cells. The indicated cell lines were seeded on glass coverslips and stained with an antibody against the V5 tag to visualise the V5-tagged recombinant TDP-43 proteins and with DAPI to visualise the nucleus. Representative images are shown. Z-stack images are covering 14 consecutive optical slices of 0.35 µm thick. Scale bar: 20 µm

### Supp Figure 7

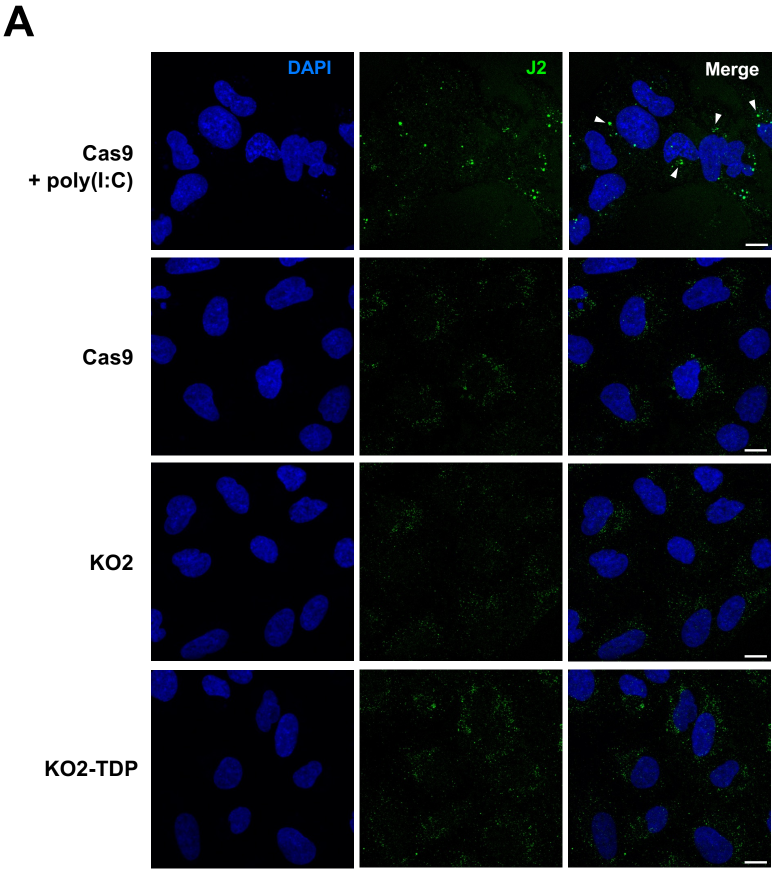


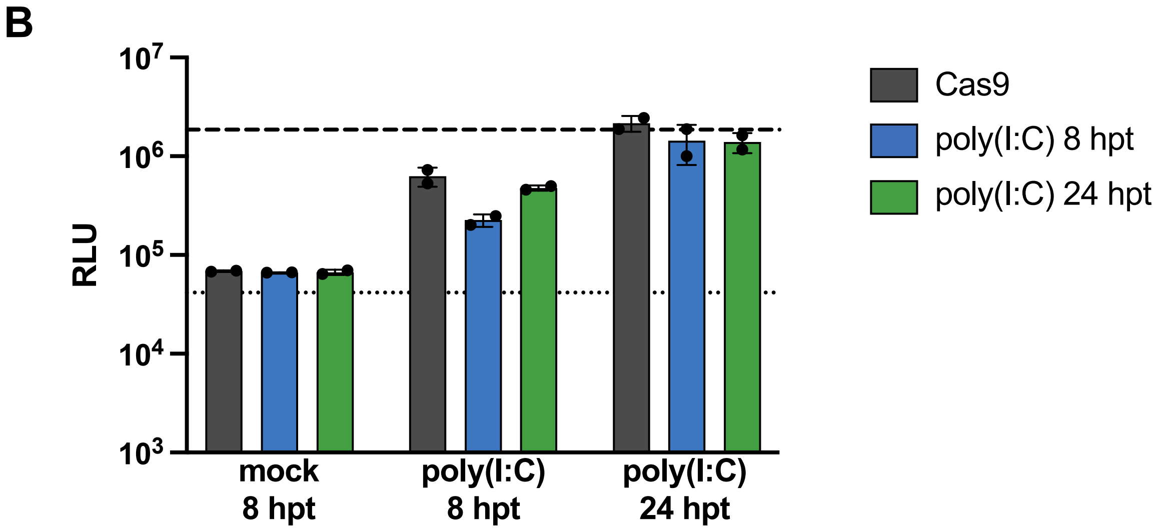


**Supp Figure 7. Effect of TDP-43 knock-out on innate immunity.**
**A.** Effect of TDP-43 knock-out on the endogenous levels of double-stranded RNAs. The Cas9, KO2 and KO2-TDP cell lines were seeded on glass coverslips and stained with an antibody specific for double-stranded RNAs (J2) and with DAPI to visualise the nuclei. Cas9 transfected with poly(I:C) were used as a positive control. Representative images are shown. White arrowheads indicate J2-positive foci. Z-stack images are covering 14 consecutive optical slices of a depth of 0.30 µm. Scale bar: 10 µm
**B.** Effect of TDP-43 knock-out on the IFN response induced by poly(I:C) treatment. The Cas9, KO2 and KO2-TDP cell lines were transfected with 500 ng of poly(I:C) or mock-transfected. The supernatants were collected at 8 and 24 hpt and transferred onto STING-37 reporter cells that express the Firefly luciferase under the control of an IFN-inducible promoter. The STING-37 cells were lysed after 24 h of incubation and the luciferase activities were measured. The dotted and dashed lines represent the background signal (STING-37 cells incubated with standard medium) and the positive control (STING cells transfected with 100 ng of poly(I:C)), respectively. The data shown are the mean ± SD of technical duplicates.
